# Single-chip End-to-End Ingestible Electronics for Gut Neurotransmitter Sensing

**DOI:** 10.64898/2026.03.28.715054

**Authors:** Angsagan Abdigazy, Mohammad Shafiqul Islam, Sandra Lara Galindo, Md Farhad Hassan, Xiang Zhang, Wooseong Choi, Mark McHugh, Sudipta Saha, Hossein Hashemi, Dong Song, Yasser Khan

## Abstract

Neurotransmitters in the gut play a vital role in human health and neuroscience, and their real-time monitoring is essential for understanding underlying physiological mechanisms. However, bioelectronic systems capable of measuring neurotransmitters *in vivo* at the anatomical site of interest remain underdeveloped and largely depend on bulky, off-the-shelf electronic components, thereby constraining the development of systems that are both practical and minimally invasive. Here, we report a miniature ingestible pill that is capable of real-time *in vivo* sensing of two key neurotransmitters: serotonin (5-HT) and dopamine (DA). The system incorporates a fully printed three-electrode-based electrochemical sensor for neurotransmitter sensing and a custom application-specific integrated circuit (ASIC) that integrates all major functional blocks on a single chip, enabling a platform for fully wireless monitoring of gut neurotransmitters. The pill, measuring 5.8 mm in diameter and 19 mm in length, supports multiple electrochemical sensing techniques, including amperometry and voltammetry, with only 42 *μ*A of average current consumption. We demonstrate the ingestible platform through *in vivo* studies in rat animal models, enabling real-time monitoring of gut neurotransmitters.

## 1 Introduction

Neurotransmitters in both the central and peripheral nervous systems play essential roles in maintaining homeostasis by regulating emotion, cognition, motor control, and gastrointestinal (GI) function. Beyond their classical roles in synaptic signaling, they also influence gut motility, nutrient absorption, immune activity, and microbiome composition. Dysregulation of key neurotransmitters such as dopamine (DA) and serotonin (5-HT) has been linked to conditions including inflammatory bowel disease (IBD), irritable bowel syndrome (IBS), and Parkinson’s disease.^[1]^ The gut–brain axis further highlights the connection between emotional state and GI function, with measurable differences in plasma neurotransmitter levels and gut microbiome profiles between IBS patients and healthy individuals, underscoring the interplay among neurotransmitter signaling, microbial composition, and psychological factors.^[2]^

Within the central nervous system (CNS), DA and 5-HT regulate cognition, emotion, and memory, and their imbalance contributes to neurological and psychiatric conditions such as depression, addiction, schizophrenia, and Parkinson’s disease.^[3]^ Outside the CNS, approximately 95% of serotonin is produced by enterochromaffin cells in the gut, where it modulates GI motility, secretion, and gut–brain communication.^[4]^ Dopamine is produced both centrally and by enteric neurons, influencing motor control, reward pathways, and GI function.^[5]^

Therefore, developing an end-to-end system capable of wirelessly measuring gut neurotransmitters in real-time is critical to elucidate the mechanisms underlying their physiological functions. In the research community, the development of sensing systems has been asymmetric. Most prior work has focused on creating robust sensors with high sensitivity and selectivity, typically validated on the bench, *in vivo* or *ex vivo* using commercial readout electronics.^[3, 6–12]^ Complete wireless ingestible systems with both sensing and communication capabilities remain rare, and are often built from off-the-shelf components that are not optimized for the given application in terms of area or power consumption.^[13–17]^ Most neurotransmitter sensors used in existing ingestible pills rely on complex fabrication processes and employ aptamers or enzymes as the sensing elements. Although aptamer- and enzyme-based sensors demonstrate high selectivity under controlled benchtop conditions, they often lack the robustness required for reliable *in vivo* operation.^[18–20]^ During the sensing data collection, many of these devices rely on Bluetooth Low Energy (BLE) for wireless communication, which supports high data rates and small antennas but suffers from high tissue attenuation. This limits communication range, increases power consumption, and can lead to unreliable connections.^[21, 22]^ Device size is another critical consideration. Larger capsules can house more components but increase the risk of GI tract retention, while smaller capsules reduce this risk and would improve patient acceptance of using these devices.^[21]^ Achieving miniaturization requires reducing the sensor size along with integrating primary electronics such as analog front end (AFE), analog-to-digital converter (ADC), radiofrequency (RF) transceiver (TRX), and secondary components such as power management unit (PMU), clock and serial communication, on a single chip to reduce capsule volume. Additionally, reducing the capsule size also means the battery used to power the electronics will be smaller, necessitating the on-chip circuitry to be ultra-low power along with being area-efficient.

In this work, we present a single-chip end-to-end ingestible pill design for neurotransmitter sensing (Fig. 1a-b), including sensor development, custom ASIC design, and wireless data management system (Fig. 1c). Our fully wireless, ultra-low-power ingestible pill is capable of real-time sensing of DA and 5-HT using a 3-electrode electrochemical sensor (Fig. 1d). To the best of our knowledge, it is the smallest ingestible pill presented in the literature, measuring 5.8 mm in diameter and 19 mm in length (Fig. 1e-g). We developed a single-chip ASIC that integrates all essential circuitry, including an AFE for sensor current-to-voltage conversion, an ADC for digitization, and an RF transmitter (TX) for wireless data communication. Unlike many existing systems, it can operate in the 402–405 MHz Medical Implant Communication System (MICS) or 433 MHz ISM (Industrial, Scientific, and Medical) bands, which provides a more favorable trade-off for ingestible devices by reducing tissue attenuation while lowering antenna area and power requirements.^[21]^ The ASIC also incorporates secondary blocks such as PMU and serial communication, enabling a complete miniaturized system-on-chip solution. In our prior work, we demonstrated a single-chip ingestible device for electrochemical sensing, which at the time was the smallest ingestible platform for such measurements.^[23]^ However, that prototype relied solely on chronoamperometry, which is well-suited for detecting biomarkers at high concentrations. For neurotransmitters present in the human body at tens to hundreds of nanomolar levels, chronoam-perometry lacks sufficient sensitivity and fails to provide accurate detection.^[3]^ To overcome this limitation, the device presented here supports a multiplexed detection of 5-HT and DA using differential pulse voltammetry (DPV) over a 10 nM to 1 *μ*M range, and chronoamperometry in 1 *μ*M to 100 *μ*M range. Finally, we demonstrate wireless detection of 5-HT in two different *in vivo* studies designed to elevate 5-HT levels: chocolate administration and dextran sulfate sodium (DSS)-induced colitis. In both cases, the changes in 5-HT concentration were monitored in real time using our ingestible pill platform (Fig. 1h).

**Figure 1.**
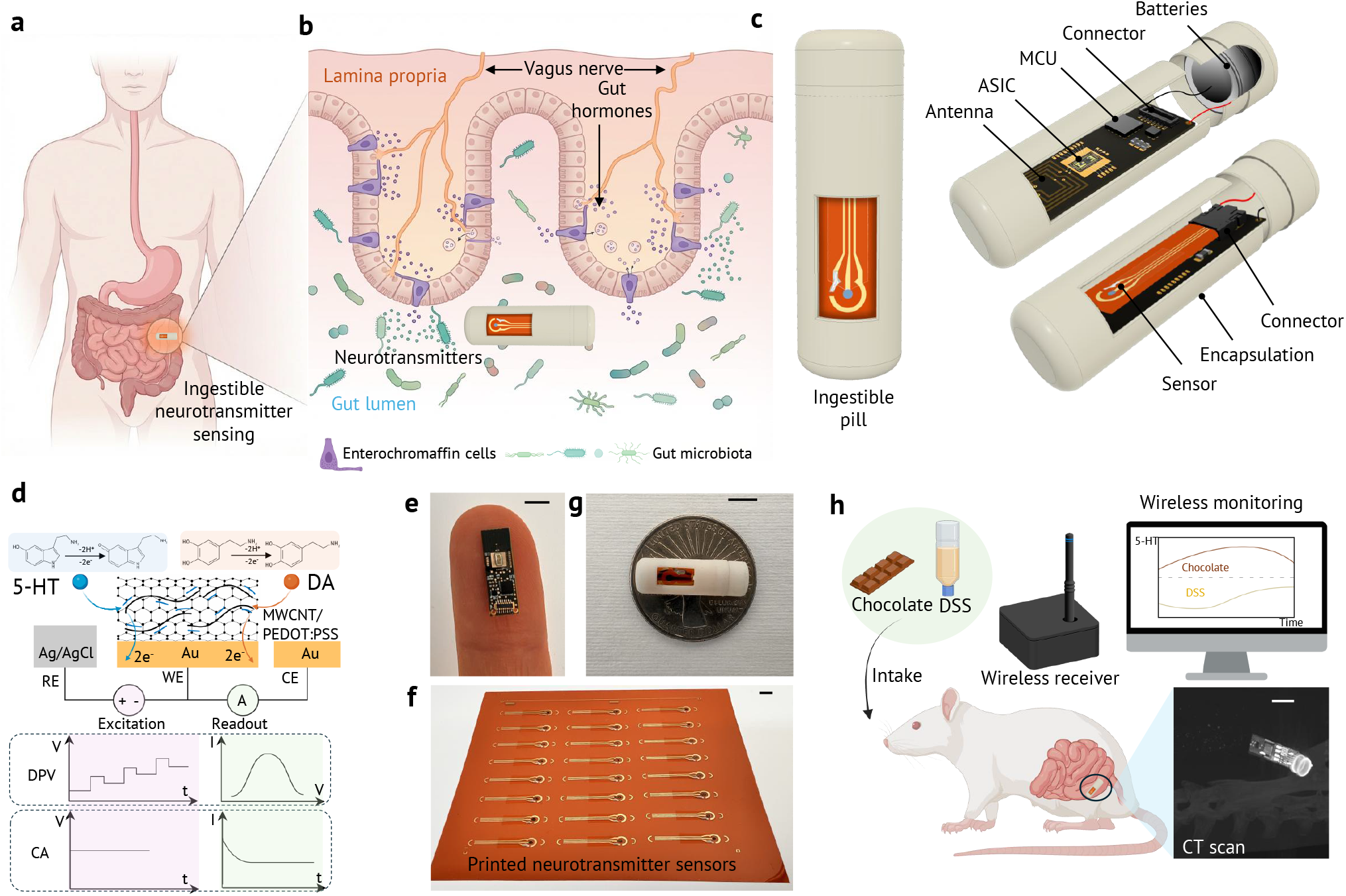
Single-chip end-to-end ingestible pill design for neurotransmitter sensing in the gut. **(a)** Schematic illustration of the human GI tract, indicating the colon as the target site for neurotransmitter sensing using the ingestible pill. **(b)** A zoomed-in view of the pill within the GI tract, along with a conceptual diagram showing vagus nerve signaling and neurotransmitter release in the gut. The developed ingestible pill will be located in the lumen for local detection of neurotransmitters. **(c)** Schematic of the fully assembled ingestible pill, showing different components, including the sensor, custom ASIC, antenna, batteries, connector, MCU, and encapsulation. **(d)** Sensing mechanism of the MWCNT/PEDOT:PSS modified electrode for neurotransmitter detection, illustrating the 3-electrode configuration (WE, RE, CE) and corresponding electrochemical techniques (DPV and CA), as well as the associated oxidation reaction for DA and 5-HT. Photographs of the **(e)** PCB with all components, **(f)** printed neurotransmitter sensor array, and **(g)** fully assembled pill displayed on a U.S.A. quarter coin. All scale bars are 5 mm. **(h)** Schematic of the assembled ingestible pill and data collection system for detecting 5-HT i*n vivo* during chocolate and DSS experiments using a rat model.

## 2 Results

Existing fabrication techniques for ingestible neurotransmitter sensors rely heavily on microfabrication, which makes the manufacturing process complex.^[19]^ In addition, aptamer-based sensing typically limits the sensor to detecting a single neurotransmitter, and many aptamer-based sensors suffer from fouling in the gastrointestinal environment, as well as complex aptamer processing steps.^[16, 24]^ Here, we present a fully printed flexible three-electrode sensor capable of detecting both DA and 5-HT across a broad concentration range, making it suitable for reliable neurotransmitter monitoring in ingestible sensing platforms.

### 2.1 Electrochemical sensor fabrication and characterization

A 3-electrode electrochemical sensor has been developed for the selective sensing of 5-HT and DA concentrations. As shown in Fig. 2a, we utilize inkjet printing to fabricate the working electrode (WE), counter electrode (CE), and reference electrode (RE) using gold nanoparticle ink (Au) on a flexible polyimide substrate. The Au ink is cured at 210°C for 1 hour.^[25]^ For the RE functionality, we apply a direct 3D printing technique to create a silver/silver-chloride (Ag/AgCl) layer on top of the printed Au electrode, which is then cured at 130°C for 25 minutes. To functionalize the WE, we employ a mixture of multi-walled carbon nanotubes (MWCNT) and Poly(2,3-dihydrothieno-1,4-dioxin)-poly(styrenesulfonate) (PEDOT:PSS), which is cured at 130°C for 20 minutes. Finally, to protect the connector from liquid exposure, we use dielectric encapsulation. The dielectric material is inkjet printed and then cured using UV light. Fig. 2b shows the scanning electron microscope (SEM) images of printed Au electrodes and the functionalized WE surface with MWCNT/PEDOT:PSS mixture.

**Figure 2.**
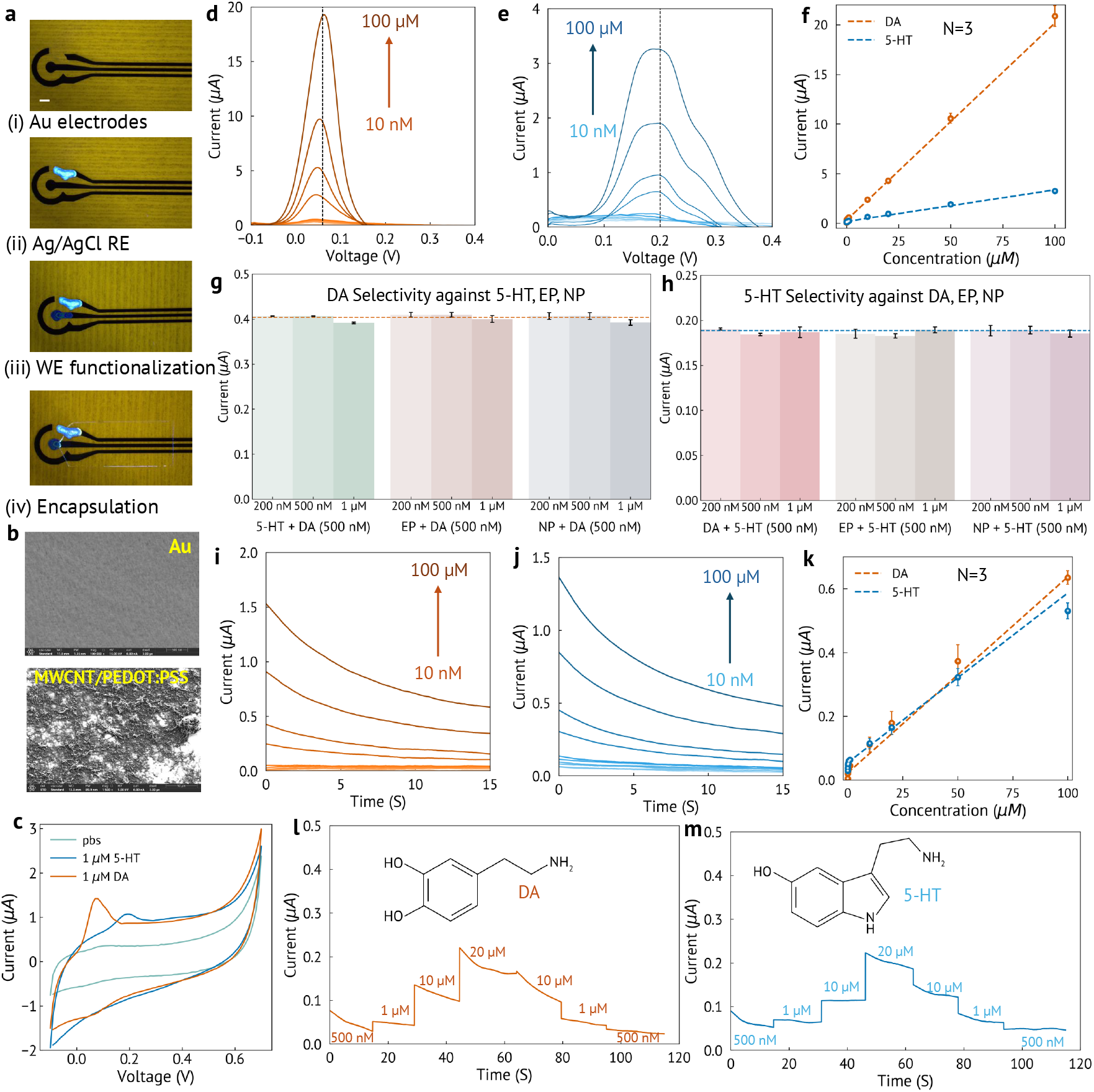
3-electrode neurotransmitter sensor design and characterization. **(a)** Micrographs illustrating the fabrication process of the sensor, including (i) inkjet-printed gold (Au) for working (WE), Reference (RE), and Counter (CE) electrodes, (ii) direct 3D-printed silver/silver-chloride (Ag/AgCl) RE, (iii) WE functionalization with MWCNT/PEDOT:PSS, and (iv) dielectric encapsulation of the connectors. Scale bar: 1 mm. **(b)** SEM images of the WE surface before (Au) and after (MWCNT/PEDOT:PSS) functionalization. **(c)** Cyclic Voltammetry in PBS, 1 *μ*M dopamin (DA) and 1 *μ*M serotonin (5-HT) solution. The sensor shows oxidation peaks at 0.06 V and 0.2 V for DA and 5-HT, respectively. **(d–e)** Differential Pulse Voltammetry (DPV) responses for DA and 5-HT (10 nM–100 *μ*M) showing peaks at 0.06 V and 0.2 V, respectively. **(f)** DPV calibration curves for DA and 5-HT based on the PBS subtracted peak DPV current values. Error bars represent the s.d. of the mean from 3 sensors. **(g)** DA (500 nM) selectivity against 5-HT, EP, and NP (200 nM–1 *μ*M) showing minimal interference. **(h)** 5-HT (500 nM) selectivity against DA, EP, and NP (200 nM–1 *μ*M) showing minimal interference. **(i–j)** Chronoamperometry (CA) responses for DA and 5-HT at potentials 0.06 V and 0.2 V, respectively. **(k)** CA calibration curves for DA and 5-HT (10 nM–100 *μ*M) based on the PBS subtracted average current values of the final 5 s of CA currents. Error bars represent the s.d. of the mean from 3 sensors. **(l-m)** Reversibility test of DA and 5-HT sensing using CA, indicating that the sensors can effectively capture variations in neurotransmitter concentrations, ranging from low to high and back again.

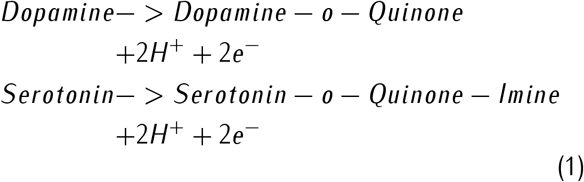

The MWCNT/PEDOT:PSS modified WE surface offers a highly conductive and electrocatalytic interface for the oxidation reactions of DA and 5-HT. DA, a catecholamine, contains two adjacent hydroxyl groups (a catechol),^[26]^ while 5-HT is a monoamine characterized by an indole ring and a hydroxyl group.^[27]^ Both neurotransmitters undergo a two-electron, two-proton oxidation to form their respective quinone and quinone-imine derivatives, as shown in equation 1. The negatively charged PSS facilitates the oxidation reaction of the neurotransmitters, while PEDOT ensures efficient charge transport to the electrode. The incorporation of MWCNTs provides a high surface area and reduces the oxidation potential (Fig. S1). The unique oxidation sites of DA (catechol –OH groups) and 5-HT (phenolic –OH group) enable selective detection with the developed sensor.

Benchtop characterization of the developed sensor is performed using a PalmSens potentiostat, beginning with CV. 10 cycles of CV in phosphate-buffered saline (PBS) are performed before each experiment to allow the sensors to stabilize (Fig. S2). As shown in Fig. 2c, the CV in PBS does not exhibit any oxidation peaks, while DA and 5-HT demonstrate oxidation peaks at 0.06 V and 0.2 V, respectively. DPV enhances measurement sensitivity by using the difference in current before and at the end of each applied potential pulse, thereby minimizing capacitive (non-faradaic) current contributions. Fig. 2d (Fig. 2e) displays the DPV currents for DA (5-HT) at concentrations ranging from 10 nM to 100 *μ*M. To eliminate the background current, the PBS DPV current is subtracted from the DPV current of the DA and 5-HT solutions. The PBS subtracted peak current values at 0.06 V and 0.2 V are used to establish the DPV calibration curves for DA and 5-HT. Fig. 2f presents the linear calibration curve for DPV detection of DA and 5-HT, demonstrating sensitivity from 200 nM to 100 *μ*M. The error bar represents the s.d. of the mean from 3 sensors. This calibration curve allows for the determination of unknown concentrations of DA and 5-HT by analyzing the PBS subtracted peak current at 0.06 V and 0.2 V, respectively. To assess the selectivity of our developed sensors for DA and 5-HT detection, the influence of other neurotransmitters is evaluated. Each solution contains a fixed concentration of primary neurotransmitter and a varying concentration of one of the interfering neurotransmitters. While keeping the primary neurotransmitter DA concentration fixed at 500 nM, the interfering neurotransmitters 5-HT, epinephrine (EP), and norepinephrine (NP) are varied from 200 nM to 1 *μ*M. Fig. 2g (Fig. S3) shows that the PBS-subtracted peak current at 0.06 V for these cocktail solutions remains at around the peak current value of 500 nM DA from the DPV calibration curve of 2f, confirming the DA selectivity against 5-HT, EP, and NP. Similarly, Fig. 2h (Fig. S4) shows the 5-HT (500 nM) selectivity against DA, EP, and NP with the concentration ranging from 200 nM to 1 *μ*M by utilizing PBS subtracted DPV peak current at 0.2 V. The sensor’s repeatability is demonstrated by performing 4 consecutive measurements of DA and 5-HT (Fig. S5). The selectivity and repeatability assessments validate the detection of DA and 5-HT, demonstrating robustness against potential interferences present within the solution.

The chronomamperometric (CA) measurements of DA and 5-HT are performed with oxidation potentials set at 0.06 V and 0.2 V, respectively. The CA current is recorded for 15 seconds to allow it to stabilize, and the PBS CA current is subtracted for background correction. Fig. 2i (Fig. 2j) illustrates the PBS subtracted CA current responses for DA (5-HT) concentrations ranging from 10 nM to 100 *μ*M. The average current value from the final 5 seconds is utilized to establish the CA calibration curves. Fig. 2k displays the linear calibration curves for both DA and 5-HT, with sensitivities ranging from 200 nM to 100 *μ*M. Finally, a repeatability test is conducted by varying the neurotransmitter concentration from low to high and back again. Fig. 2l (Fig. 2m) shows the PBS-subtracted CA current response as the concentration of DA (5-HT) fluctuates from 500 nM to 20 *μ*M and then back to 500 nM, confirming the sensor’s capability to detect continuously varying concentrations of the neurotransmitter solutions. Following the sensor development, we designed the readout electronics to enable a fully integrated ingestible pill system (Fig. S6).

### 2.2 Electronic circuit design and characterization

Fig. 3a shows a system-level overview of the proposed device. It consists of a printed circuit board (PCB) that integrates all electronic components, an electrochemical sensor, and coin-cell batteries. Fig. 3b shows a single rigid FR-4 PCB with dimensions of 5 mm × 15 mm × 1.2 mm, which integrates a custom application-specific integrated circuit (ASIC), measuring 1 mm × 2 mm × 300 *μ*m and taped out in TSMC 180 nm CMOS process, directly wire-bonded onto PCB pads and encapsulated in epoxy, a loop coil serving as the antenna for the on-chip RF transmitter, a microcontroller unit (MCU), an FPC connector for interfacing with a 3-electrode electrochemical sensor, a board-to-board (B2B) connector for initial MCU programming, capacitors, and two contact pads for battery connections. Fig. S7 shows the schematic and layout of the PCB, and Supplementary Table 1 lists all electronic components. Fig. 3c presents the die micrograph of the ASIC, which integrates an analog front-end (AFE) for 3-electrode electrochemical sensing, an analog-to-digital converter (ADC) to digitize the AFE output, a digital-to-analog converter (DAC) to generate control voltages for the AFE and ADC, a radio-frequency (RF) transmitter for wireless data transmission to an external reader, a power management unit (PMU) consisting of a bandgap voltage reference (BGR) and three low-dropout regulators (LDOs) supplying the analog, digital, and RF blocks, and a scan chain used for communicating and programming on-chip modules via the MCU.^[23]^ Two silver-oxide batteries connected in series were used to power the device, providing a total supply voltage of 3.1 V and a combined capacity of 7.5 mAh. A measured battery lifetime of 4 days was achieved, which could be further extended by a factor of three by reducing the duty cycle through less frequent measurements and increased sleep duration (Fig. S8).

**Figure 3.**
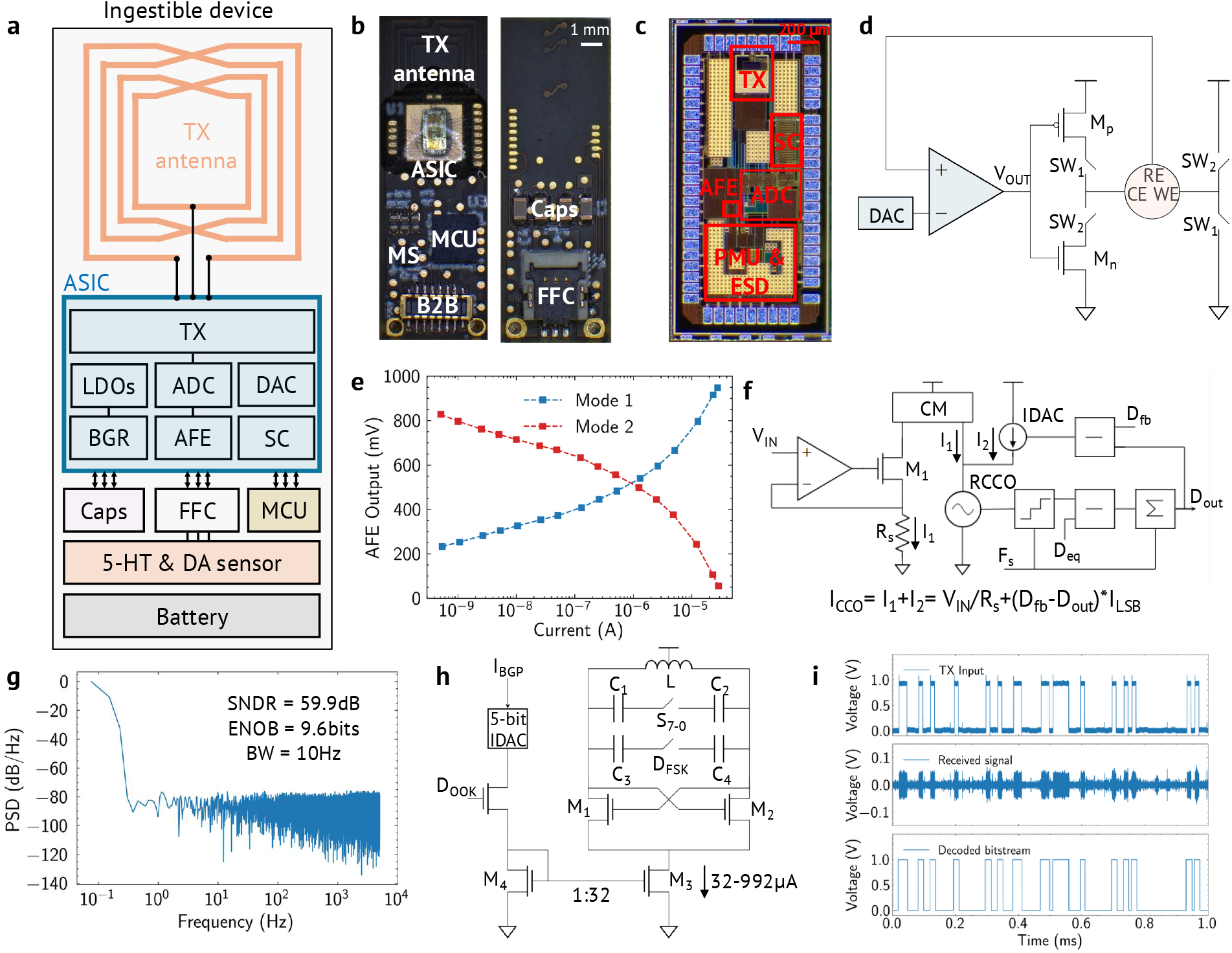
System-level overview and ASIC characterization. **(a)** System-level overview of the device, comprising a printed circuit board (PCB) integrating all electronics, a 3-electrode sensor, and two silver-oxide batteries. **(b)** Images of the 4-layer PCB from top (left) and bottom (right) sides. Scale bar: 1 mm. **(c)** Die micrograph of the ASIC fabricated in the TSMC 180 nm process node, measuring 1 mm × 2 mm × 300 *μ*m. Scale bar: 200 *μ*m. **(d)** Schematic of the AFE, consisting of a rail-to-rail input folded-cascode operational transconductance amplifier (OTA), a common-source stage directly interfacing the sensor, and a resistive voltage DAC for sensor biasing. **(e)** Measured output voltage of the AFE at sensor currents ranging from sub-nA to 29 *μ*A for both oxidation and reduction reactions. **(f)** Block diagram of the closed-loop VCO-based ADC, used to convert the AFE output voltage to a digital bitstream. **(g)** Measured power spectral density (PSD) of the ADC output. The ADC achieves a signal-to-noise and distortion ratio (SNDR) of 59.9 dB, corresponding to 9.6 bits of effective number of bits (ENOB). **(h)** Schematic of the LC VCO-based TX, which sends the ADC bitstream to the external receiver. **(i)** Measured transmitted and received transient plots of a PRBS sequence using on-off keying (OOK) modulation.

The analog front-end (AFE) used for 3-electrode electrochemical sensing is shown in Fig. 3d. It interfaces directly with the electrochemical sensor and enables measurement of both oxidation and reduction currents, supporting amperometric and voltammetric operation.^[28]^ The polarity between the working electrode (WE) and reference electrode (RE) is configurable through on-chip switches (SW_1_, SW_2_), allowing flexible biasing depending on the sensing modality. The RE potential is regulated through a closed-loop architecture using a 7-bit voltage digital-to-analog converter (DAC) that provides the bias input to the OTA. This programmability enables both static biasing for chronoamperometry and time-varying voltage profiles required for techniques such as cyclic voltammetry (CV), differential pulse voltammetry (DPV), and square-wave voltammetry (SWV), which are generated through the on-chip scan chain. The sensor current (*I*_*sens*_) is converted to a voltage (*V*_*out*_) at the OTA output through the feedback loop. As electrochemical measurements are dominated by low-frequency signals, flicker noise becomes the primary limitation. To address this, the AFE employs a chopper-stabilized folded-cascode OTA with a rail-to-rail input stage, enabling both low-noise operation and a wide input range for the WE–RE potential difference. The chopping frequency is set to 20 kHz and generated on-chip. As a result, the integrated inputreferred noise over 1 mHz-100 Hz remains below 100 pA_*rms*_ (Fig. S9a). The AFE is designed for ultra-low-power operation, with the OTA, DAC, and chopping circuitry consuming 800, 100, and 100 nW, respectively. By utilizing a singleamplifier topology, the design minimizes both area (80 *μ*m *×* 40 *μ*m) and power consumption compared to conventional potentiostat implementations. To evaluate performance, the AFE was characterized using a standard 3-electrode equivalent circuit model implemented with discrete components. As shown in Fig. 3e, the system supports a bidirectional current range from 0.5 nA to 29 *μ*A (*±*29 *μ*A). The stability of the feedback loop across this range is confirmed by phase margin simulations (Fig. S9b).

The analog-to-digital conversion is implemented using a closed-loop, time-based architecture built around a current-controlled ring oscillator (Fig. 3f). In this approach, the input voltage (*V*_*in*_) is first translated into a current that modulates the oscillation frequency of a five-stage ring oscillator. The resulting frequency-encoded signal is digitized by counting oscillator transitions and processing the accumulated counts through a digital feedback loop. The loop operates by continuously comparing the oscillator output to a reference corresponding to its equilibrium frequency *f*_0_, and correcting deviations through a feedback current generated by a binary-weighted current DAC (IDAC). This establishes a self-regulating system in which the oscillator is driven toward its steady-state frequency, while the digital output (*D*_*out*_) tracks the input signal. The feedback term (*D*_*fb*_) enables offset adjustment to compensate for process, voltage, and temperature (PVT) variations. The IDAC is referenced to the on-chip bandgap and defines the least significant bit current (*I*_*LSB*_). This time-domain conversion scheme enables a compact and energy-efficient implementation, as it relies primarily on digital operations and avoids large passive components. The ADC supports an input range from 100 mV to 900 mV while maintaining a small footprint (360 *μ*m *×* 300 *μ*m). As the electrochemical signals of interest vary slowly relative to the sampling rate, the input can be treated as quasi-static during conversion. Accordingly, the ADC performance is evaluated based on its noise characteristics under constant input conditions. The signal-to-noise ratio (SNR) and effective number of bits (ENOB) are extracted from the power spectral density (PSD) of the output bitstream, normalized to full scale.^[29]^ Fig. 3g shows the measured PSD for a DC input of *V*_*in*_ = 500 mV at a sampling rate of 8 kS/s, and the ADC achieves an ENOB of 9.6 bits with a conversion time of 100 ms, while consuming 1.2 *μ*W of power.

Wireless data transmission is achieved using an LC VCO-based transmitter that operates in the 402–405 MHz Medical Implant Communication System (MICS) and the 433 MHz ISM bands (Fig. 3h). These frequency ranges provide a favorable trade-off for ingestible devices by balancing tissue attenuation, antenna size, and achievable data rates.^[21]^ The transmitter employs a cross-coupled LC oscillator whose frequency is digitally tunable through an 8-bit capacitive DAC (CDAC), enabling operation across a wide range from 350 to 450 MHz. An off-chip loop coil implemented on the PCB serves a dual role as both the inductive element of the LC tank and the radiating antenna, reducing system complexity and footprint. The coil consists of four turns with outer dimensions of 3.6 mm × 3.6 mm, trace width and spacing of 6 mil, achieving an inductance of 64 nH and a quality factor of 80 at 400 MHz based on electromagnetic simulations (Fig. S10). The oscillator bias current is programmable through a current DAC, allowing dynamic control of transmit power. The transmitter operates with a minimum startup current of 64 *μ*A and supports tuning up to 992 *μ*A, corresponding to an output power range from -44 dBm to -25 dBm (Fig. S11a). Both on–off keying (OOK) and frequency-shift keying (FSK) modulation schemes are supported to accommodate different communication requirements (Fig. S11b). Measured transient responses for OOK transmission at 80 kb/s are shown in Fig. 3i, while the corresponding eye diagram demonstrates reliable data recovery with a bit error rate below 10^−4^ (Fig. S11c). During experiments, a compact off-the-shelf 433 MHz receiver module (MAX1473) with a whip antenna was used to capture the transmitted signal (Fig. 1h). At the minimum bias current, the VCO exhibits a phase noise of -110 dBc/Hz at a 300 kHz offset (Fig. S11d). The total on-chip area of the transmitter is 250 *μ*m × 250 *μ*m.

The on-chip power management unit (PMU) generates regulated supply rails using a bandgap reference (BGR) and multiple low-dropout regulators (LDOs). Independent domains are provided for the analog, digital, and radio-frequency blocks to ensure stable operation across the system.^[30]^ The LDO outputs are programmable from 0.9 V to 1.25 V (nominally 1 V) and support load currents of up to 2 mA. External decoupling capacitance of 10 *μ*F is used at the regulator outputs to maintain stability. The PMU is optimized for low-power operation, consuming less than 2 *μ*W in total, and enables the entire system to be powered from a single 1.55 V battery.

### 2.3 Validation on the bench and *in vivo* animal models

After separate characterization of the sensor and electronics, the complete system was assembled for further validation. The sensor was connected to the proposed electronic system via an FFC connector (see Fig. 3b) and characterized on the bench to verify compatibility with the ASIC. In the first experiment, the DAC in the ASIC’s AFE was continuously programmed by the MCU to perform differential DPV with parameters matched to those of a commercial potentiostat used for initial characterization of sensors. The device was exposed to solutions containing DA and 5-HT at concentrations ranging from 10 nM to 1 *μ*M to evaluate system performance and generate calibration curves. The AFE output was recorded and post-processed in MATLAB to extract the sensor current as a function of the applied potential, as shown in Fig. 4a–b. The current values measured in PBS solution were subtracted for background correction. From these waveforms, the peak currents at 0.12 V for DA and 0.22 V for 5-HT were used to construct the corresponding calibration curves, which are also shown in Fig. 4a–b. In the next experiment, the MCU programmed the DAC in the AFE to apply fixed potentials of 0.12 V and 0.22 V for DA and 5-HT, respectively. CA is performed using solutions containing DA and 5-HT at concentrations ranging from 10 nM to 100 *μ*M, with the corresponding results shown in Fig. 4c-d. This is a one-time calibration required for the new electronics developed with the ASIC, and there is no need to repeat as long as the electronics and sensor design are same.

**Figure 4.**
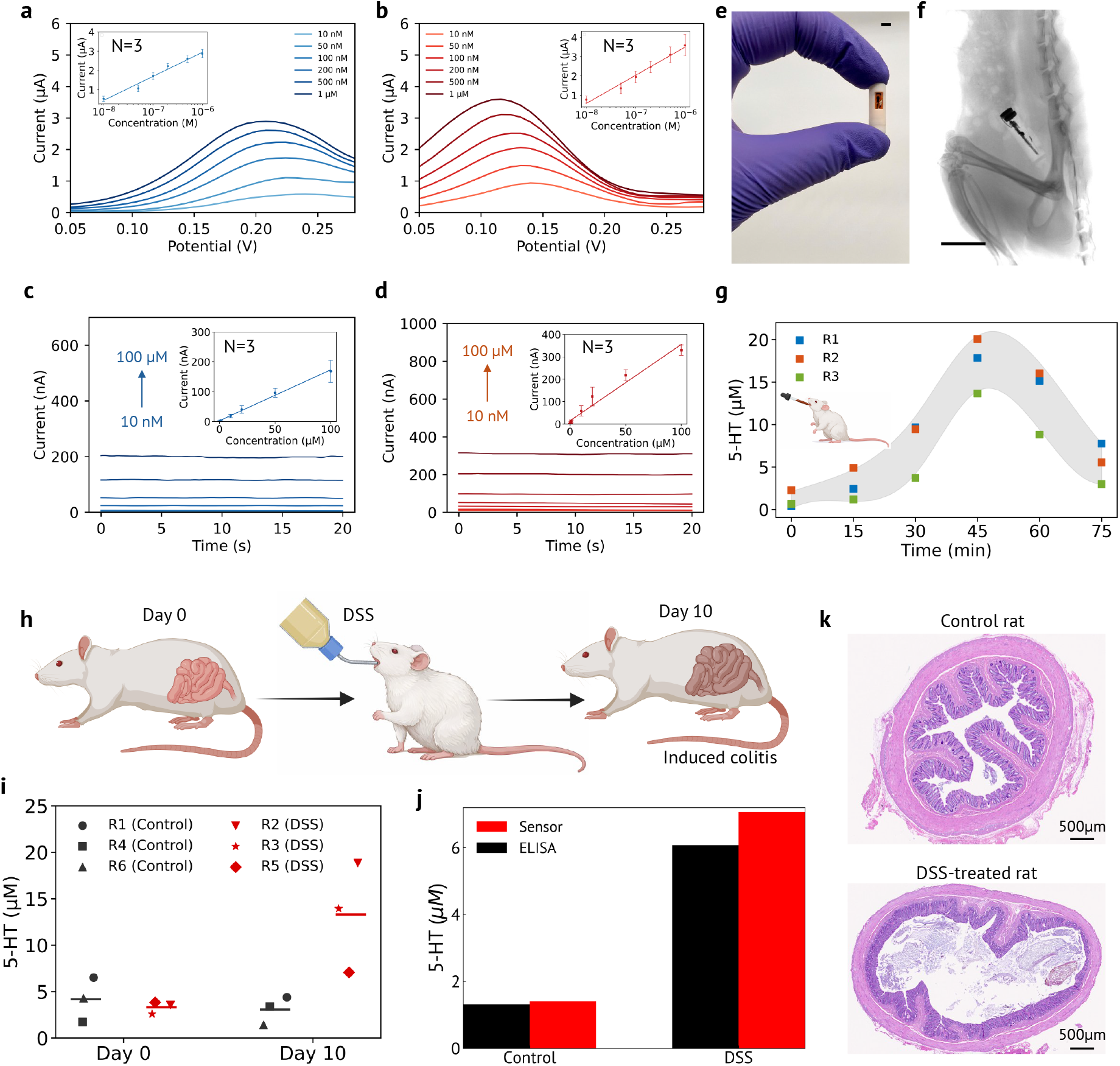
Bench-top and *in vivo* validation results. **(a)** Bench-top measurements of 5-HT using DPV. Inset shows the linear calibartion curve for 5-HT (N = 3). **(b)** Bench-top measurements of DA using DPV. Inset shows the linear calibartion curve for DA (N = 3). **(c-d)** Bench-top measurements of 5-HT and DA using chronoamperometry at 0.22 V and 0.12 V, respectively.. Inset shows the linear calibartion curves (N = 3). **(e)** Photograph of the proposed ingestible device. Scale bar: 5 mm. **(f)** X-ray scan of a rat with the pill inserted into the colon showing the pill position during the *in vivo* experiments. Scale bar: 2 cm. **(g)** *In vivo* monitoring of 5-HT over time in the rat colon after chocolate syrup administration. Experiments are repeated in 3 rats. **(h-i)** *In vivo* measurements of 5-HT levels in the rat colon during DSS-induced acute colitis development. A total of 6 rats are used: 3 control and 3 DSS-treated. **(j)** ELISA measurements of colonic 5-HT levels in one control (R6) and one DSS-treated (R5) rat after 10 days. **(k)** Histology images showing H&E-stained colon tissues of control and DSS-treated rats after 10 days. The colonic tissue of DSS-treated rats exhibited clearly visible damage to the GI lining.

After successful bench-top validation of the complete system, we proceeded with *in vivo* experiments in the colon of the living rat models (Supplementary Video 1). Fig. 4e shows the complete ingestible pill. Fig. 4f shows the location of the pill after *in vivo* experiments through X-ray imaging (Supplementary Video 2 shows the CT-scan). In the first experiment, the rats are fasted for 3 hours prior to the procedure. Chocolate syrup was then administered orally using a syringe with a small feeding tube. The rats are immediately anesthetized with 1-2% isoflurane, and the lubricated pill is inserted into the distal colon via the anus. The 5-HT concentration is monitored over the following 75 minutes with the first measurement (t = 0 min) right after the rats are anesthetized. The experiment is repeated in three rats. As shown in Fig. 4g, similar 5-HT dynamics are observed across all animals: colonic 5-HT gradually increases over time, peaks within 35–55 minutes, and then gradually decreases over the next 20–40 minutes. In the second *in vivo* experiment, two groups of rats are studied: a control group and an acute colitis group. Acute colitis are induced by administering dextran sulfate sodium (DSS) in the animals’ drinking water^[31]^ (Fig. 4h). Baseline 5-HT levels in the colon of all rats are first recorded on Day 0. After 10 days of DSS exposure, the measurements are repeated, with the results shown in Fig. 4i. As observed, the 5-HT levels in the DSS group increase significantly over the 10-day period, while there is no significant change in the 5-HT concentrations of the control group. To quantify, the R2 (DSS) 5-HT concentration increased from 3.55 *μ*M (day 0) to 18.85 *μ*M (day 10). These elevated 5-HT concentrations are due to the fact that DSS exposure damages the colonic epithelium and is associated with an increased density of 5-HT producing enterochromaffin (EC) cells in the colon.^[3]^ Following the *in vivo* experiments, distal colon tissues harvested from the six rats are utilized for histology (all six rats) and the gold standard Enzyme-Linked Immunosorbent Assay (ELISA) (two rats, one from each group) analyses, with the respective results shown in Fig. 4j-k and Fig. S14-15. Fig. 4j shows strong agreement between *in vivo* measurements obtained using our ingestible pill and gold-standard ELISA analysis, validating our ingestible pill platform for reliable nuerotransmitter sensing. As it can be seen in Fig. 4k, the colonic tissue of DSS-treated rats exhibited clearly visible damage of GI lining due to induced colitis.

## 3 Discussion

In this work, we present an end-to-end ingestible platform for real-time *in vivo* neurotransmitter sensing. Supplementary Table 2 compares the proposed device with previously reported ingestible electrochemical sensing systems. To the best of our knowledge, this system is the most compact device reported to date that enables real-time electrochemical sensing with *in vivo* validation, achieving at least a 1.9× reduction in size and an 8.3× reduction in average power consumption compared to the state of the art.^[16]^ Miniaturization is critical not only for broader clinical adoption, but also for fundamental research applications. In particular, studies of gastrointestinal disease and the gut–brain axis often rely on small animal models such as rats, rabbits, and minipigs, where device size is a key constraint. Continued reductions in size and power consumption are therefore essential for enabling these devices as practical research tools.

In our *in vivo* colon experiments, we observed slightly higher absolute serotonin levels than those reported in prior studies.^[3, 16]^ This difference may be attributed to our experimental approach, in which the rat colon was not cleaned prior to device placement in order to better reflect realistic physiological conditions. While patients undergoing capsule endoscopy or other ingestible diagnostics typically undergo bowel preparation and fasting, residual fluids and microbial content remain present in the gastrointestinal tract, which can influence measured neurotransmitter levels.

In the future, the AFE design can be further improved and optimized to enhance overall system performance. A next-generation ASIC can incorporate a receiver for realtime remote configuration, along with on-chip memory and digital filters for local data storage and processing. In addition, integrating a precise clock with a finite-state machine (FSM) enables autonomous execution of voltammetry techniques, eliminating the need for a microcontroller. From a sensing perspective, the platform can be extended to support multiplexed detection of additional biomarkers, including pH, glucose, and other neurotransmitters. Progression toward clinical translation can be supported through larger-animal *in vivo* studies, such as in porcine models.

## Methods

### Sensor fabrication

A 15 *μ*m thick polyimide flexible substrate (Xenomax, Nagase) was used for the sensor fabrication. The WE, RE, and CE were patterned by inkjet printing Au nanoparticle ink (Harima Chemicals), followed by curing at 210°C for 1 h. To complete the RE fabrication, Ag/AgCl ink (Insulectro) was printed on top of the Au layer and cured at 130°C for 25 min. The Ag/AgCl layer was fabricated using direct 3D writing (Voltera) with a print speed of 200 mm min^-1^, nozzle diameter of 225 *μ*m, and vacuum power of 60%. The print height was set to 150 *μ*m with a trace spacing of 180 *μ*m. The dispense pressure and relief pressure were 600 and 30, respectively, and the dispenser preheat temperature was maintained at 35°C. The WE was subsequently functionalized with a MWCNT/PEDOT:PSS composite. Initially, 0.1 wt% of MWCNTs(Sigma Aldrich) were dispersed in 1mL of DI water containing 0.1 vol% of Triton X-100 (Sigma Aldrich) and sonicated to obtain a homogeneous suspension. This dispersion was then added to 1mL of 0.8 wt% aqueous PEDOT:PSS (Sigma Aldrich) treated with 5 vol% DMSO (Sigma Aldrich). The final mixer was sonicated for 5 min prior to deposition. A volume of 0.4 *μ*L of the final composite mixer was drop-cast onto each WE and kept at room temperature overnight. It was cured at 130°C for 20min. Finally, dielectric encapsulation was inkjet printed over the connector region to prevent the direct contact with the liquid. A SU-8-based dielectric ink (Kayaku Advanced Materials) was diluted in a 2:1 ratio with CPG thinner (Kayaku Advanced Materials) prior to printing. The printed encapsulation layer was exposed to a UV dose of 35 mJ/cm^2^ for 25 min, followed by a soft bake at 110°C for 2 min. All the inkjet printing processes were carried out using a FUJIFILM Dimatix DMP 2850 printer equipped with a 2.4 pL drop volume printhead and a drop spacing of 20 *μ*m. For Au nanoparticle printing, a jetting potential of 30 V and a substrate temperature of 40°C were used, while dielectric encapsulation printing was done with 33 V of jetting potential and 45°C of substrate temperature.

### Sensor characterization

Benchtop characterization of the neurotransmitter sensors was performed using a PalmSens potentiostat. Prior to measurements, the sensors were stabilized by conducting 10 cycles of CV in PBS. For DPV and CA measurements of DA and 5-HT, a pretreatment step was implemented to enhance sensitivity. Specifically, a 900 s deposition step was applied at a low potential (0 V for DA and -0.05 V for 5-HT) while the solution was stirred magnetically to promote the accumulation of analytes on the electrode surface. For DPV measurements, a step potential of 0.002 V, pulse potential of 0.075 V, and pulse time of 0.1 s were used. For CA measurement, constant DC potentials of 0.06 V and 0.2 V were used for DA and 5-HT, respectively. A sampling interval of 0.02 s was used. For all DPV and CA current measurements, the PBS current was subtracted for background correction.

### Device assembly

The printed circuit board (PCB) presented in this work was fabricated on a standard four-layer FR4 substrate with dimensions of 15 mm × 5 mm × 0.4 mm. The PCB incorporates a micro-controller (STMicroelectronics, STM32L011E4Y6TR), a board-to-board connector (Molex, 5050701222), a flexible printed circuit (FPC) connector (Kyocera, 046277003001883+), four 22 *μ*F capacitors (Kyocera, 04026D226MAT2A), and three 2.7 kΩ resistors (Yageo, RC0201JR-072K7L), all soldered on both sides of the board. The custom ASIC, fabricated by Muse Semiconductor using the TSMC 180 nm CMOS process, was wire-bonded directly onto the PCB pads by ProtoConnect LLC and encapsulated with clear epoxy to protect the interconnections. The loop antenna for the radiofrequency (RF) transmitter (TX) was designed using copper traces on the PCB itself. Two 1.55 V silver oxide batteries (SR416SW, 4.8 mm × 1.6 mm, 7.5 mAh each) were connected in series and soldered to the PCB pads using thin copper wires. The device enclosure was 3D printed using a Form 3B+ printer (Formlabs) with BioMed Clear resin (FLBMCL01), a biocompatible medical-grade material (see Fig. S12). A dedicated opening was incorporated into the shell design to expose the sensor interface to the GI environment. A biocompatible silicone elastomer (NuSil MED-4011) was used to seal the gap between the sensor and encapsulation opening.

### ASIC characterization

The ASIC was first tested in a benchtop environment. To characterize the analog front end (AFE) and extract its calibration curve, relating output voltage to sensor current, we implemented a sensor model consisting of resistors and capacitors on a separate test PCB and breadboard.^[32]^ This model was connected to the ASIC test board. The model parameters, particularly the resistance between the working and reference electrodes, were precisely varied, and the corresponding AFE output voltage was recorded. To minimize loading effects, the AFE output was connected to an external voltage buffer before being measured using a mixed-signal oscilloscope (Keysight MSOS104A). To characterize the analog-to-digital converter (ADC), we used a Digilent Analog Discovery 3, which integrates a waveform generator, power supply, oscilloscope, and logic analyzer. Various voltage waveforms were applied to the ADC input, and the serialized digital outputs were captured using the logic analyzer. The recorded bit streams were exported and post-processed in MATLAB to verify linearity and resolution. For radiofrequency (RF) transmitter (TX) characterization, an external PRBS-7 sequence generated by the Analog Discovery 3 was used as the digital input at different data rates. The external reader setup consisted of a whip antenna (ANT-418-CW-HWR-SMA) placed 1 m from the chip, a low-noise amplifier (ZRL-700+), and either an oscilloscope or a spectrum analyzer, depending on the measurement type. After benchtop characterizations, an off-the-shelf handheld receiver module (MAX1473) with the same whip antenna was used to receive the data from the device. Finally, to measure the output voltages from the bandgap voltage reference (BGR) and low-dropout regulators (LDOs), these nodes were first connected to an external voltage buffer on the test board, and the buffered outputs were recorded using the oscilloscope of the Analog Discovery 3.

### Animal studies

#### Experiments involving animal subjects

All sensing procedures for rats were achieved in conformity with protocols certified by the Institutional Animal Care and Use Committee (IACUC) at the University of Southern California. Approximately at the time of the experiments, 2 month old male Sprague Dawley rats were used and weighed 420 grams on average. Six rats were used for the in-vivo experiments. Rats were single housed and kept in pairs during the duration of the experiments. They were kept at 22°C, 12-h light/dark cycle, and with access to food and water unless otherwise noted.

#### Serotonin sensing in the colon of living rats

Prior to the colon sensing in the rats, the animals were fasted for 3 hours in advance to the experiment. The 100*μ*L of Hershey’s chocolate syrup was administered orally via using a syringe with small feeding tube. They were then anaesthetized with 1–2% isoflurane (variation in concentration are due to the slight difference in the weight of the rats) and placed in a stereotaxic apparatus while resting on a heating pad to maintain the animal’s temperature. The device was placed into plastic medical grade tubing as the shuttle with medical lubricant to ease the insertion into the colon via anus. Chronoamperometry with a potential of 220 mV was used to monitor the serotonin dynamics over time. The measurements were performed at the distal colon region 5 cm from the anus. Following the experiment, animals were returned to their cages to have a full recovery.

#### Colitis inflammation model

Rats were induced into acute colitis by administering DSS in drinking water 2.5% wt./vol, for 10 days from day 0 changing the water every two days. During and after DSS treatment, animals were monitored for signs of pain, altered feeding habits or weight loss to ensure that their nutritional needs are met and there is no notable distress.^[31]^ The serotonin in the colon was measured in living anaesthetized rats, similar to the procedure described in the previous section.

#### Histology analysis

After completion of all *in vivo* measurements, the animals were euthanized and the distal colon was harvested from each of the six rats for histology analysis. The extracted colon tissue was kept in neutral buffer with 10% formalin for 24 hours, then embedded in paraffin. The tissue section with a thickness of 5 *μ*m was prepared and stained with H&E staining.

#### ELISA analysis

After completion of *in vivo* measurements, animals were euthanized, and the distal colon was harvested from R6 of the control group and R5 of the DSS-treated group for ELISA analysis. The extracted colon samples were snap-frozen at -80°C until further processing. For protein extraction, tissues were placed on a cold surface with the mucosal side facing upward. A cell scraper was held at a 45° angle to gently remove the superficial epithelium along with some goblet cells. The collected material was transferred into cold 1× PBS buffer and washed twice with cold 1× PBS. Subsequently, 200 *μ*L 1X Lysus buffer was added to the cell pellet. The samples were mixed and incubated on ice for 30 minutes, followed by centrifugation at 12,000 g for 10 minutes at 4°C. The supernatant was then transferred to a new tube. The concentrations of DA and 5-HT were quantified using commercially available ELISA kits (5-HT: RayBiotech; DA: AssayGenie).

## Lead contact

Further information and requests for resources should be directed to the lead contact, Yasser Khan.

## Data and code availability

Any additional data included in this manuscript and the supplemental information are available from the lead contact upon request.

## Acknowledgments

Yasser Khan acknowledges support from the Packard Fellowship and the Air Force Office of Scientific Research Young Investigator Program.

## Author contributions

A.A., M.S.I., and Y.K. conceived the project. A.A. designed and characterized the ASIC, PCB, and complete electronic system. M.S.I. designed, fabricated, and characterized the sensors. S.L.G. and X.Z. developed the animal protocol. M.F.H. designed the pill casing, and S.S. simulated the Au-CNT surface. A.A., M.S.I., S.L.G., X.Z., W.C., and M.M. carried out *in vivo* studies. H.H. supervised the ASIC design. D.S. supervised the *in vivo* studies. A.A., M.S.I., S.L.G., and Y.K. wrote the manuscript. All authors reviewed and have given approval to the final version of the manuscript.

## Declaration of generative AI and AI-assisted technologies in the writing process

During the preparation of this manuscript, the authors used Chat-GPT to check for grammatical errors in the writing. After using this tool, the authors reviewed and edited the content as needed and take full responsibility for the content of the publication.

## References

[1] Mittal, R. et al. Neurotransmitters: The critical modulators regulating gut–brain axis. Journal of cellular physiology 232, 2359–2372 (2017).

[2] Barandouzi, Z. A. et al. Associations of neurotransmitters and the gut microbiome with emotional distress in mixed type of irritable bowel syndrome. Scientific Reports 12, 1648 (2022).

[3] Li, J. et al. A tissue-like neurotransmitter sensor for the brain and gut. Nature 606, 94–101 (2022).

[4] Liu, H. N., Nakamura, M. & Kawashima, H. New role of the serotonin as a biomarker of gut–brain interaction. Life 14, 1280 (2024).

[5] Mhanna, A. et al. The correlation between gut microbiota and both neurotransmitters and mental disorders: A narrative review. Medicine 103, e37114 (2024).

[6] Orzari, L. O., de Freitas, R. C., de Araujo Andreotti, I. A., Gatti, A. & Janegitz, B. C. A novel disposable self-adhesive inked paper device for electrochemical sensing of dopamine and serotonin neurotransmitters and biosensing of glucose. Biosensors and Bioelectronics 138, 111310 (2019).

[7] Cernat, A., Ştefan, G., Tertis, M., Cristea, C. & Simon, I. An overview of the detection of serotonin and dopamine with graphene-based sensors. Bioelectrochemistry 136, 107620 (2020).

[8] Kumar, S. et al. A machine learning approach for simultaneous electrochemical detection of dopamine and serotonin in an optimized carbon thread-based miniaturized device. IEEE Sensors Journal (2024).

[9] Ruckodanov, D. A., Maidment, N. T. & Monbouquette, H. G. Electrochemical sensing of dopamine with an implantable microelectrode array microprobe including an on-probe iridium oxide reference electrode. ACS Chemical Neuroscience (2025).

[10] Zhao, C. et al. Implantable aptamer–field-effect transistor neuroprobes for in vivo neurotransmitter monitoring. Science advances 7, eabj7422 (2021).

[11] Chapin, A. A. et al. Electrochemical measurement of serotonin by au-cnt electrodes fabricated on microporous cell culture membranes. Microsystems & Nanoengineering 6, 90 (2020).

[12] Galindo, S. L., Islam, M. S. & Khan, Y. Serotonin and dopamine sensing in the acidic gut environment. In 2025 IEEE BioSensors Conference (BioSensors), 1–4 (IEEE, 2025).

[13] Han, J. et al. Simultaneous dopamine and serotonin monitoring in freely moving crayfish using a wireless electrochemical sensing system. ACS sensors 9, 2346–2355 (2024).

[14] Straker, M. A. et al. Seropill: Novel minimally invasive ingestible capsule for serotonin sensing in the gi tract. In 2023 22nd International Conference on Solid-State Sensors, Actuators and Microsystems (Transducers), 1884–1887 (IEEE, 2023).

[15] Castagnola, E. et al. Stable in-vivo electrochemical sensing of tonic serotonin levels using pedot/cnt-coated glassy carbon flexible microelectrode arrays. Biosensors and Bioelectronics 230, 115242 (2023).

[16] Min, J. et al. Continuous biochemical profiling of the gastrointestinal tract using an integrated smart capsule. Nature Electronics 1–12 (2025).

[17] Even, A. et al. Measurements of redox balance along the gut using a miniaturized ingestible sensor. Nature Electronics 8, 856–870 (2025).

[18] Sequeira-Antunes, B. & Ferreira, H. A. Nucleic acid aptamer-based biosensors: a review. Biomedicines 11, 3201 (2023).

[19] Weltin, A., Kieninger, J. & Urban, G. A. Microfabricated, amperometric, enzyme-based biosensors for in vivo applications. Analytical and bioanalytical chemistry 408, 4503–4521 (2016).

[20] Kratschmer, C. & Levy, M. Effect of chemical modifications on aptamer stability in serum. Nucleic acid therapeutics 27, 335–344 (2017).

[21] Abdigazy, A. et al. End-to-end design of ingestible electronics. Nature Electronics 7, 102–118 (2024).

[22] Abdigazy, A. et al. 3d gas mapping in the gut with ai-enabled ingestible and wearable electronics. Cell Reports Physical Science 5 (2024).

[23] Abdigazy, A. et al. A self-orienting single-chip ingestible pill for electrochemical sensing in the gi tract. In 2024 IEEE Biomedical Circuits and Systems Conference (BioCAS), 1–5 (IEEE, 2024).

[24] Overton, S. N. et al. Serotonin sensing technologies to promote understanding of the gut–brain axis. IEEE Sensors Letters 8, 1–4 (2024).

[25] Islam, M. S. et al. Wearable organic-electrochemical-transistor-based lithium sensor for precision mental health. Device 3 (2025).

[26] Liu, X. & Liu, J. Biosensors and sensors for dopamine detection. View 2, 20200102 (2021).

[27] Coyle, V. E., Brothers, M. C., McDonald, S. & Kim, S. S. Superlative and selective sensing of serotonin in undiluted human serum using novel polystyrene sulfonate conductive polymer. ACS omega 9, 16800–16809 (2024).

[28] Tsai, J.-H.Chen, Y.-C. & Liao, Y.-T. A power-efficient bidirectional potentiostat-based readout ic for wide-range electrochemical sensing. In 2018 IEEE International Symposium on Circuits and Systems (ISCAS), 1–5 (IEEE, 2018).

[29] Sacco, E., Vergauwen, J. & Gielen, G. A 16.1-bit resolution 0.064-mm 2 compact highly digital closed-loop single-vco-based 1–1 sturdy-mash resistance-to-digital converter with high robustness in 180-nm cmos. IEEE Journal of Solid-State Circuits 55, 2456–2467 (2020).

[30] Abdigazy, A. & Monge, M. A bimodal low-power transceiver featuring a ring oscillator-based transmitter and magnetic field-based receiver for insertable smart pills. IEEE solid-state circuits letters 5, 154–157 (2022).

[31] Chassaing, B., Aitken, J. D., Malleshappa, M. & Vijay-Kumar, M. Dextran sulfate sodium (dss)-induced colitis in mice. Current protocols in immunology 104, 15–25 (2014).

[32] Ahmadi, M. M. & Jullien, G. A. Current-mirror-based potentiostats for three-electrode amperometric electrochemical sensors. IEEE Transactions on Circuits and Systems I: Regular Papers 56, 1339–1348 (2008).

