## Supplementary Information for "Single-chip End-to-End Ingestible Electronics for Gut Neurotransmitter Sensing"

**Supplementary Note 1** . . . 2

#### List of Figures

S1 Au-CNT electrostatic potential profile simulation . . . 3
S2 Cyclic Voltammetry (CV) in PBS . . . 4
S3 Selectivity of DA against 5-HT, EP and NP . . . 5
S4 Selectivity of DA against 5-HT, EP and NP . . . 6
S5 Repeatability test for DA and 5-HT detection . . . 7
S6 Fabrication and assembly of the ingestible pill . . . 8
S7 Schematic and layout of the PCB from Autodesk EAGLE PCB Designer. . . . 9
S8 Measured battery voltage over time under various duty cycles . . . 11
S9 Simulations of the AFE. . . . 12
S10 Simulations of the TX antenna . . . 13
S11 Measurements and simulations of the TX . . . 14
S12 3D printing of pill casing . . . 15
S13 Colon extraction for histology and ELISA analysis . . . 16
S14 Histology images . . . 17
S15 ELISA analysis . . . 18

### Supplementary Note 1

Au-CNT electrostatic potential profile and barrier formation analysis were performed using the DMol$^3$ module implemented in *Materials Studio*. Structural geometry optimization was carried out employing the Generalized Gradient Approximation (GGA) with the Perdew–Burke–Ernzerhof (PBE) exchange–correlation functional.
A vacuum slab of 20 Å was introduced along the surface normal (z-direction) to eliminate spurious interactions between periodic images. Structural relaxation was conducted until strict convergence criteria were satisfied. The total energy convergence threshold was set to $1.0 \times 10^{-7}$ Ha, while the force convergence criterion was fixed at $1.0 \times 10^{-4}$ Ha/Å.
To approximate the influence of the surrounding medium and screen long-range electrostatic interactions, the COSMO solvation model was employed. For simplicity and to emulate ambient conditions, water was chosen as the background dielectric medium with a dielectric constant of $\varepsilon = 78.4$ at room temperature (25 °C). In addition, the slab consisted of three layers of Au atoms, consistent with earlier study.[1]
Figure S1 presents the effective electrostatic potential profiles along the transport direction (z-axis) for (a) pristine Au, (b) Au–CNT interface with 90° orientation, and (c) Au–CNT interface with 0° orientation. The Fermi level ($E_F$) is indicated for reference.
For pristine gold (Fig. S1a), a comparatively higher potential barrier is observed near the surface region. Upon introducing CNT on top of the Au surface, the interfacial electrostatic environment is significantly modified, leading to a reduction in the effective barrier height. This reduction originates from charge redistribution and interface dipole formation at the Au–CNT junction. Importantly, the CNT orientation strongly influences the barrier characteristics. In the 0° configuration (Fig. S1c), a distinct interfacial barrier is formed between Au and CNT with a height of approximately 0.58 eV. It is consistent with the previous study.[2] In contrast, for the 90° orientation (Fig. S1b), the barrier height is significantly reduced, indicating enhanced electronic coupling and improved charge transport across the interface. Intermediate barrier heights between 0° and 90° were approximated assuming quasi-linear variation with orientation angle in order to systematically evaluate orientation-dependent tunneling probability. The electron tunneling probability was evaluated using the one-dimensional WKB (Wentzel–Kramers–Brillouin) approximation. For a rectangular barrier of height $\Phi$ and width $d$, the transmission coefficient $T(E)$ for an electron of energy $E$ is given by:

$$T(E) \approx \exp\left[-2\kappa d\right], \tag{SE1}$$

where

$$\kappa = \frac{\sqrt{2m^*(\Phi - E)}}{\hbar}. \tag{SE2}$$

Here, $m^*$ is the effective electron mass, $\hbar$ is the reduced Planck constant, $\Phi$ is the barrier height, and $E$ is the incident electron energy.

For a spatially varying barrier $V(z)$, the generalized WKB expression becomes:

$$T(E) \approx \exp\left[-2\int_{z_1}^{z_2} \frac{\sqrt{2m^*\left(V(z) - E\right)}}{\hbar}\, dz\right], \tag{SE3}$$

where $z_1$ and $z_2$ are the classical turning points defined by $V(z) = E$.

Figure S1(d) illustrates the calculated transmission coefficient $T(E)$ for various CNT orientations from 0° to 90°. As the orientation angle increases, the effective barrier height decreases, resulting in enhanced tunneling probability.

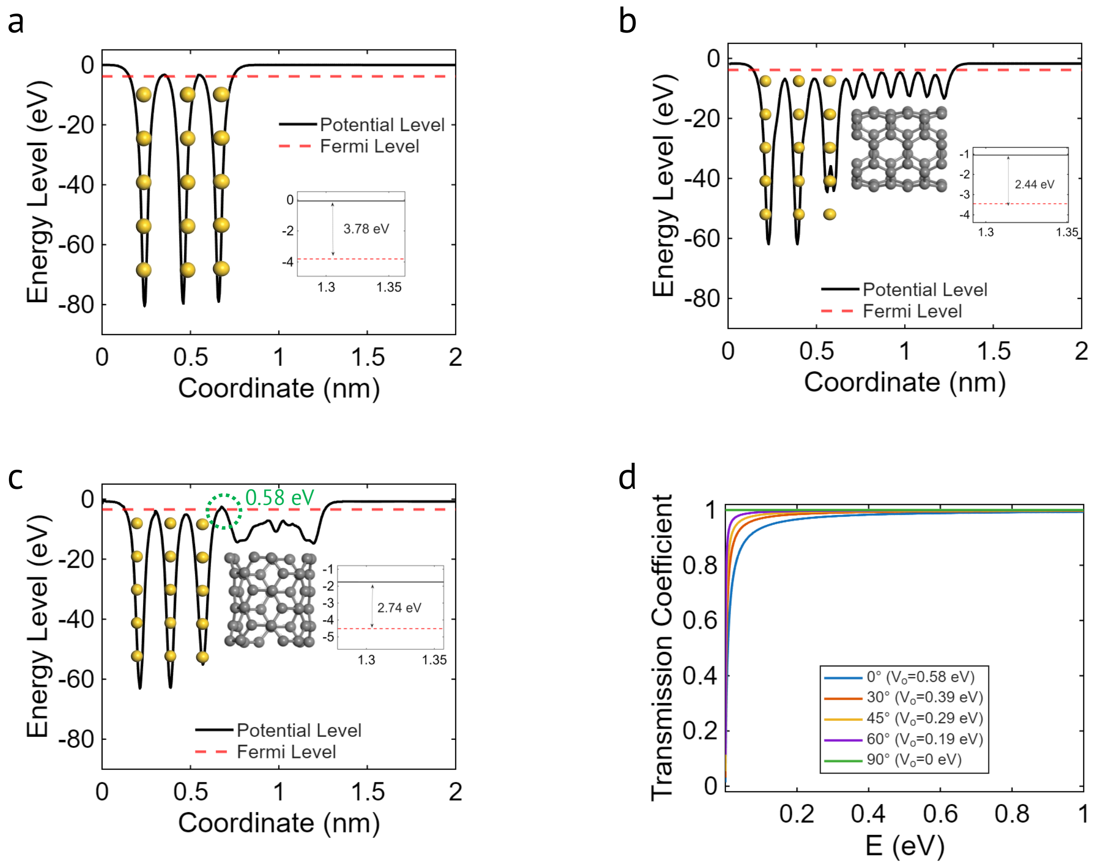

**Figure S1. Au-CNT electrostatic potential profile simulation.**
Effective electrostatic potential profiles calculated along the z-direction using DFT (GGA-PBE) for
**(a)** pristine Au surface,
**(b)** Au–CNT interface with 90° CNT orientation, and
**(c)** Au–CNT interface with 0° orientation.
A vacuum slab of 15 Å was used to prevent periodic interactions. The Fermi level ($E_F$) is indicated in each panel. The 0° configuration exhibits an interfacial barrier of approximately 0.58 eV, whereas the 90° configuration shows a significantly reduced barrier due to enhanced interfacial coupling.
**(d)** Orientation-dependent electron transmission coefficient $T(E)$ calculated using the WKB approximation, demonstrating increased tunneling probability with decreasing barrier height.

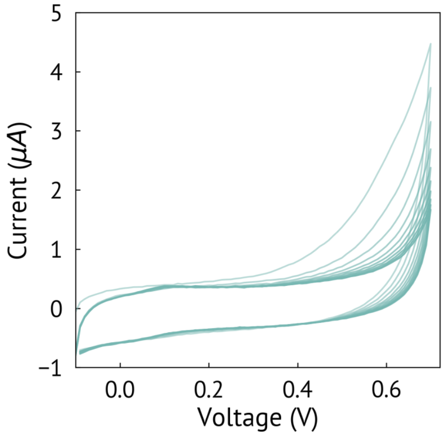

**Figure S2. Cyclic Voltammetry (CV) in PBS.**
Ten consecutive CV cycles in PBS were performed to stabilize the sensor response. The overlapping current responses indicate a steady electrochemical interface.

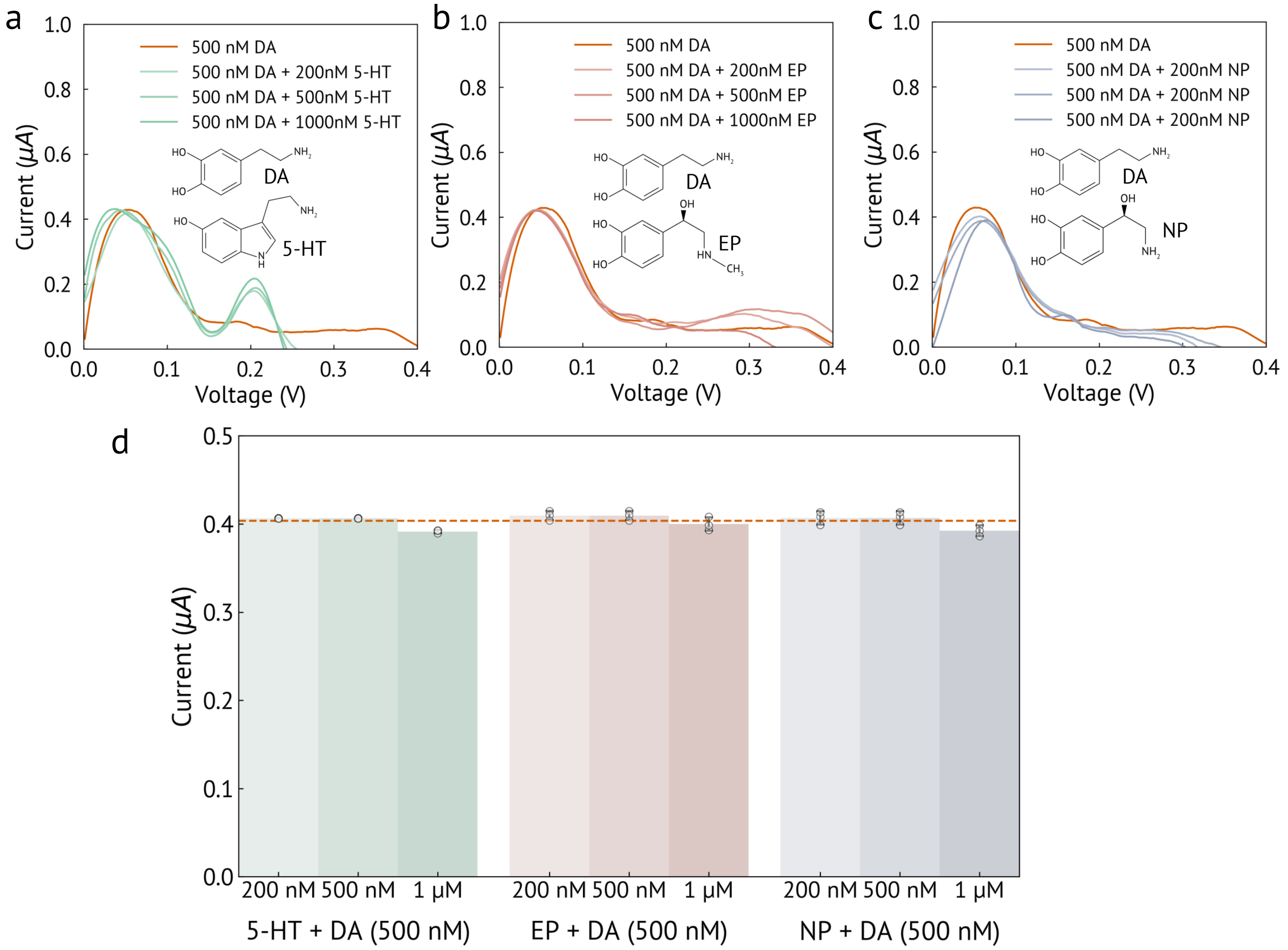

**Figure S3. Selectivity of DA against 5-HT, EP and NP.**

**(a)** DPV responses of the cocktail solution containing DA and 5-HT. The concentration of DA was fixed at 500 nM, while the 5-HT concentration was varied from 200 nM to 1 $\mu$M. The DA oxidation peak remained constant at 0.06 V, whereas the 5-HT peak current increased with increasing 5-HT concentration at 0.2 V, indicating well-resolved and independent DPV peaks.

**(b)** DPV responses of the cocktail solution containing DA and EP. The concentration of DA was fixed at 500 nM, while the EP concentration was varied from 200 nM to 1 $\mu$M. No significant change in the DA peak current or peak potential was observed, demonstrating negligible interference from EP.

**(c)** DPV responses of the cocktail solution containing DA and NP. The concentration of DA was fixed at 500 nM, and NP concentration was varied from 200 nM to 1 $\mu$M. No significant change in the DA peak current or peak potential was observed, demonstrating negligible interference from NP.

**(d)** Quantified DA DPV peak currents at 0.06 V in the presence of varying concentrations of interfering neurotransmitters (5-HT, EP, and NP). The DA peak current remained nearly constant and comparable to that observed with 500 nM DA alone, indicating minimal influence from the interfering species and confirming the sensor's high selectivity.

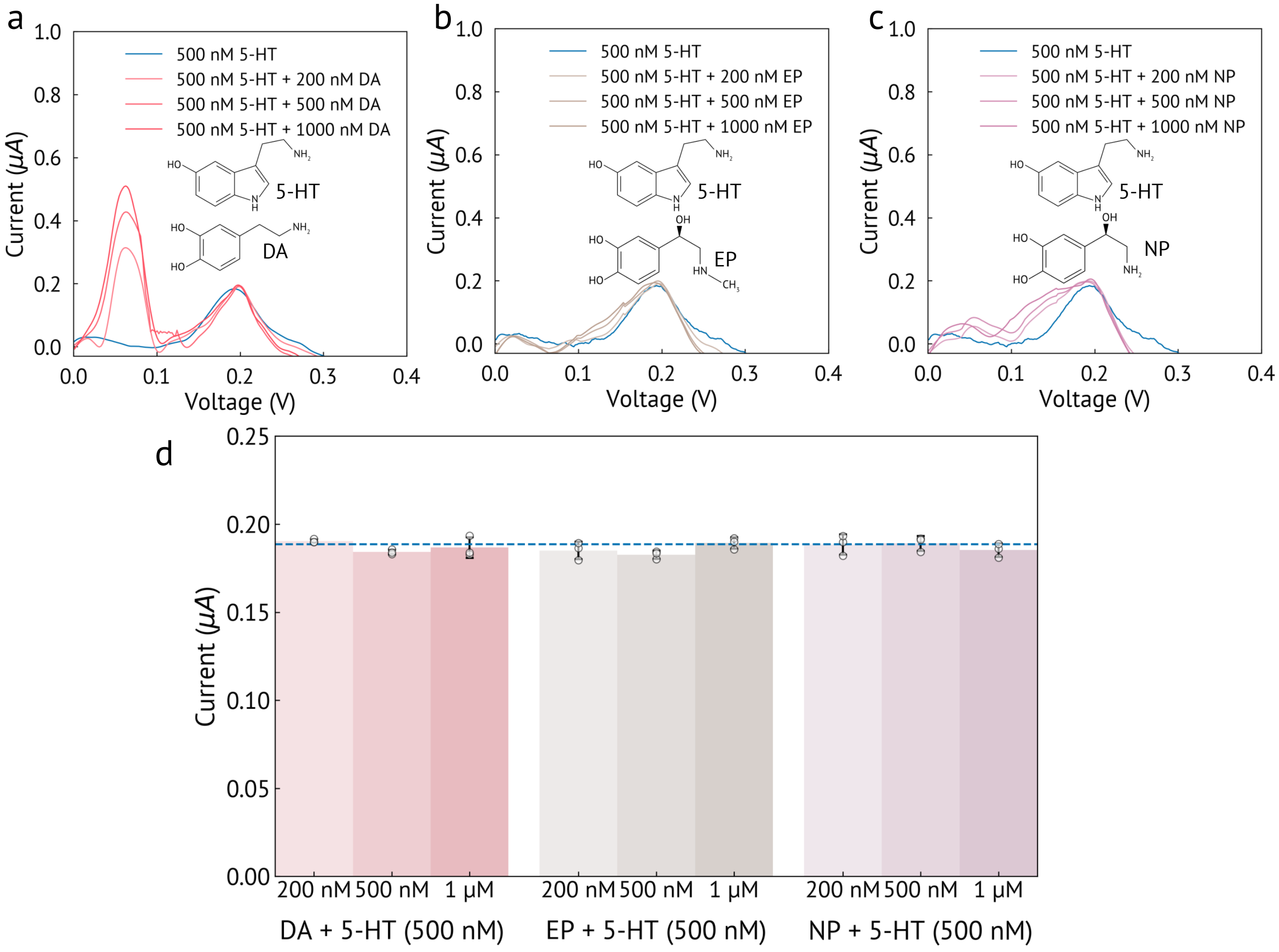

**Figure S4. Selectivity of DA against 5-HT, EP and NP.**

**(a)** DPV responses of the cocktail solution containing 5-HT and DA. The concentration of 5-HT was fixed at 500 nM, while the DA concentration was varied from 200 nM to 1 $\mu$M. The 5-HT oxidation peak remained constant at 0.2 V, whereas the DA peak current increased with increasing DA concentration at 0.06 V, indicating well-resolved and independent DPV peaks.

**(b)** DPV responses of the cocktail solution containing 5-HT and EP. The concentration of 5-HT was fixed at 500 nM, while the EP concentration was varied from 200 nM to 1 $\mu$M. No significant change in the 5-HT peak current or peak potential was observed, demonstrating negligible interference from EP.

**(c)** DPV responses of the cocktail solution containing 5-HT and NP. The concentration of 5-HT was fixed at 500 nM, while the NP concentration was varied from 200 nM to 1 $\mu$M. No significant change in the 5-HT peak current or peak potential was observed, demonstrating negligible interference from NP.

**(d)** Quantified 5-HT DPV peak currents at 0.2 V in the presence of varying concentrations of interfering neurotransmitters (DA, EP, and NP). The 5-HT peak current remained nearly constant and comparable to that observed with 500 nM 5-HT alone, indicating minimal influence from the interfering species and confirming the sensor's high selectivity.

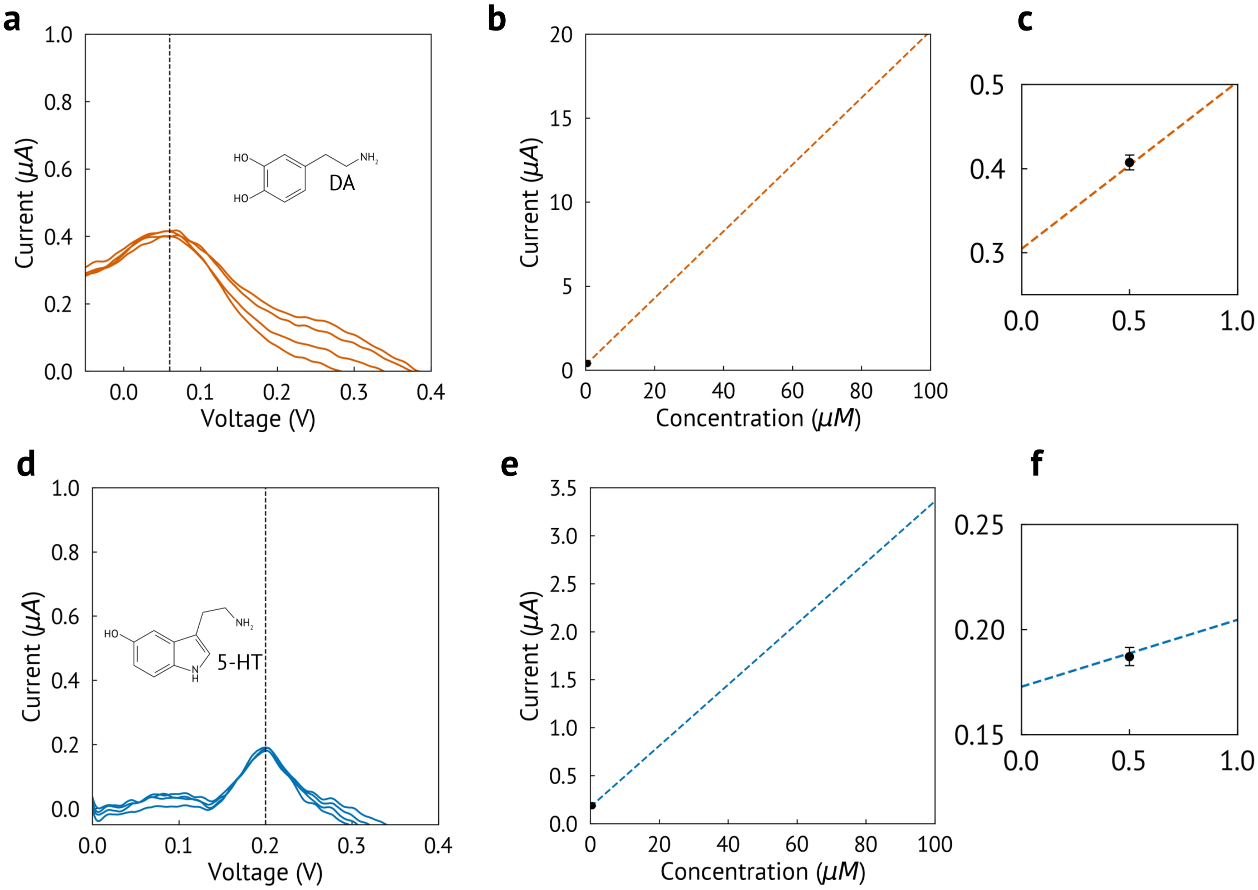

**Figure S5. Repeatability test for DA and 5-HT detection.**
**(a)** Four consecutive DPV cycles recorded for DA, demonstrating consistent peak current and peak potential, indicating good measurement repeatability.
**(b)** The peak current measured at 0.06 V for all measurements was consistent with the corresponding calibration curve, confirming the reliability and reproducibility of the sensor response for DA.
**(c)** Zoomed in view of the low concentration of S5b.
**(d)** Four consecutive DPV cycles recorded for 5-HT, demonstrating consistent peak current and peak potential, indicating good measurement repeatability.
**(e)** The peak current measured at 0.2 V for all measurements was consistent with the corresponding calibration curve, confirming the reliability and reproducibility of the sensor response 5-HT.
**(f)** Zoomed in view of the low concentration of S5e.

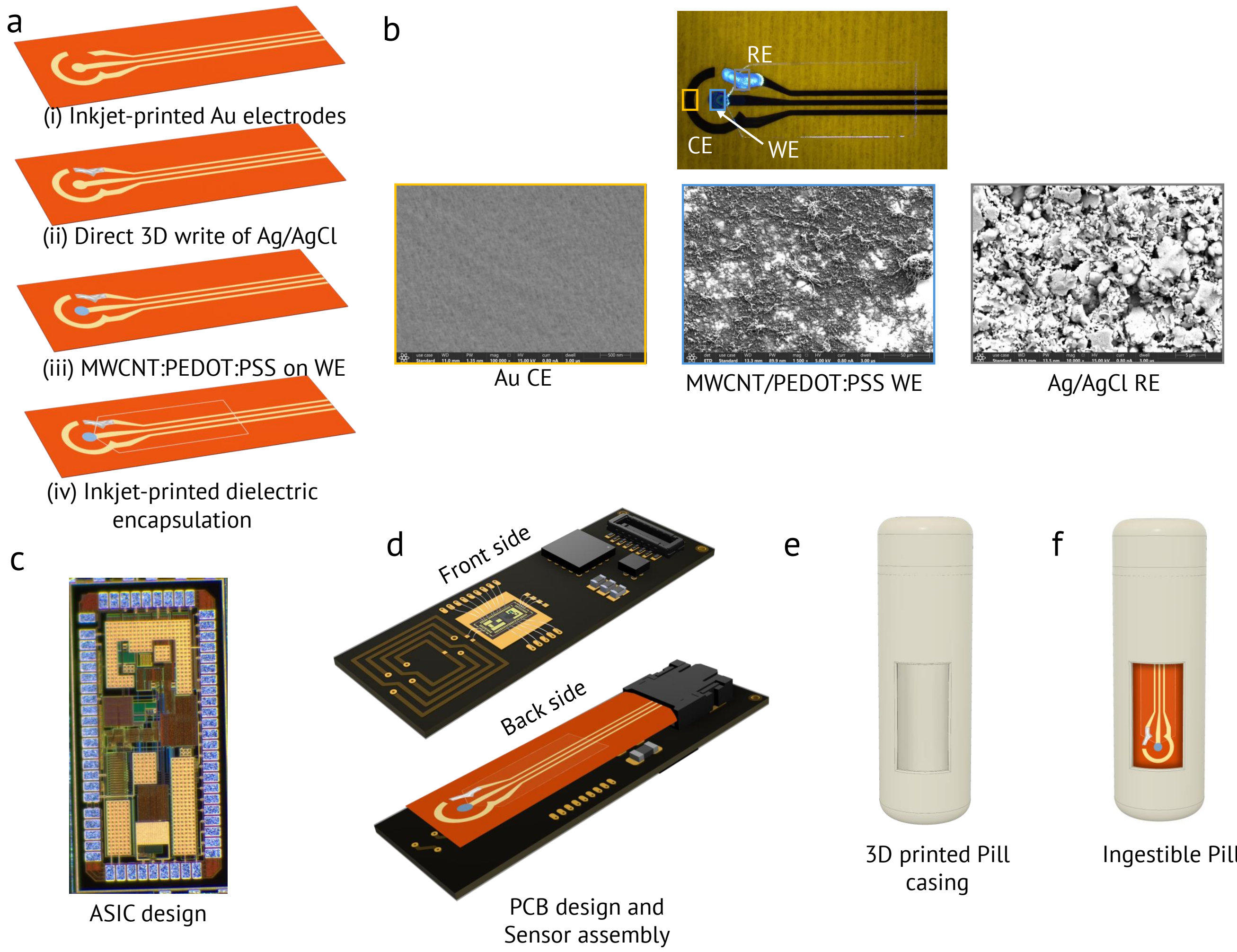

**Figure S6. Fabrication and assembly of the ingestible pill**

**(a)** Fabrication steps of the flexible neurotransmitter sensor: (i) inkjet printing of Au electrodes, (ii) direct 3D writing of Ag/AgCl on the RE, (iii) functionalization of the WE with MWCNT/PEDOT:PSS composite, and (iv) inkjet printing of dielectric encapsulation.

**(b)** Micrograph of the fabricated neurotransmitter sensor and corresponding SEM images of the Au CE, MWCNT/PEDOT:PSS modified WE, and Ag/AgCl RE.

**(c)** Micrograph of the custom designed ASIC.

**(d)** PCB design and integration of the flexible sensor with the readout electronics.

**(e)** 3D printed pill casing.

**(f)** Fully assembled ingestible pill with the integrated neurotransmitter sensor.

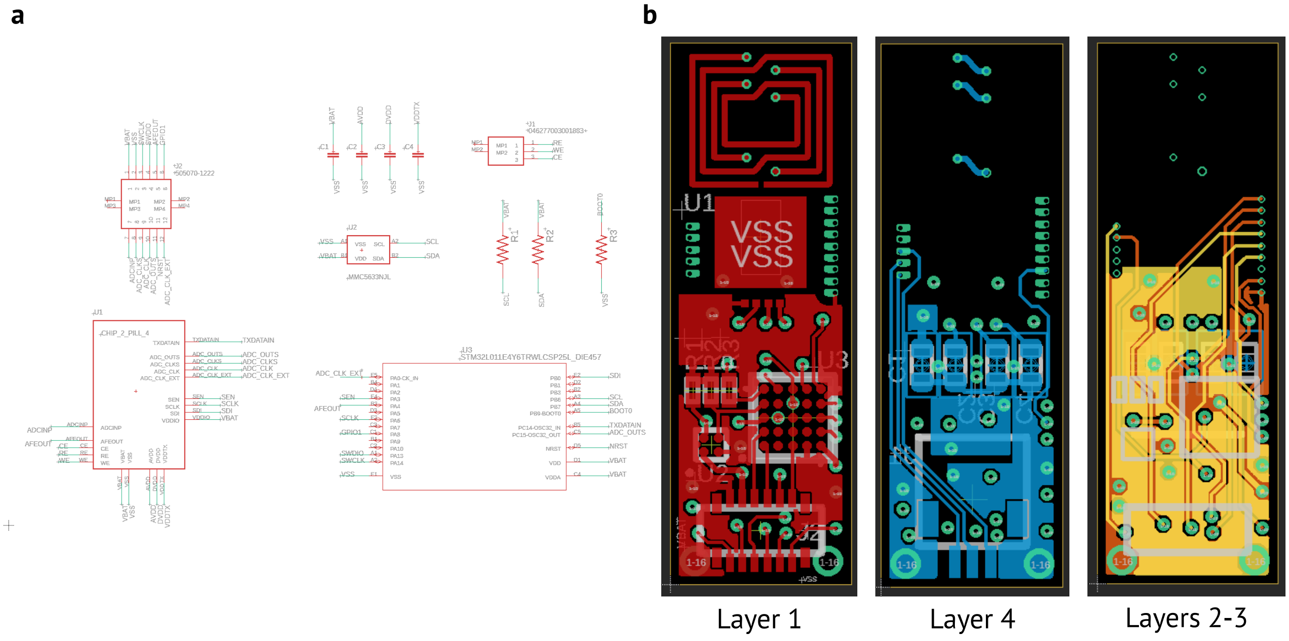

**Figure S7. Schematic and layout of the PCB from Autodesk EAGLE PCB Designer.**
**(a)** Schematic view of the PCB. U1 – ASIC; U2 – magnetic sensor for localization purposes, which was not used in this work; U3 – microcontroller unit (MCU); J1 – FFC connector for 3-electrode electrochemical sensor; J2 – B2B connector for programming and debugging purposes; R1-R2 – I2C pull-up resistors; R3 – pull-up resistor for MCU; C1-C4 – decoupling capacitors for battery and on-chip LDOs.
**(b)** Layout view of the 4-layer PCB. The top side (Layer 1) contains ASIC, TX loop antenna, MCU, magnetic sensor, resistors, and B2B connector. Layer 1 metal was primarily used for routing, and as a power plane; The bottom side (Layer 4) contains the sensor FFC connector and capacitors. Layer 4 metal was primarily used for routing, and as a ground plane; The middle layers (Layer 3-4) were used primarily as power and ground planes with some minor routing.

**Table T1.** List of PCB components

| Component | Description | Manufacturer | Part # |
|---|---|---|---|
| U1 | Custom ASIC | TSMC | N/A |
| U2 | Magnetic sensor | Memsic Inc. | MMC5633NJL |
| U3 | MCU | STMicroelectronics | STM32L011E4Y6TR |
| C1–C4 | 22$\mu$F capacitors | KYOCERA AVX | 04026D226MAT2A |
| J1 | FPC connector | KYOCERA AVX | 046277003001883+ |
| J2 | B2B connector | Molex | 5050701222 |
| R1–R3 | 2.7k$\Omega$ resistors | YAGEO | RC0201JR-072K7L |

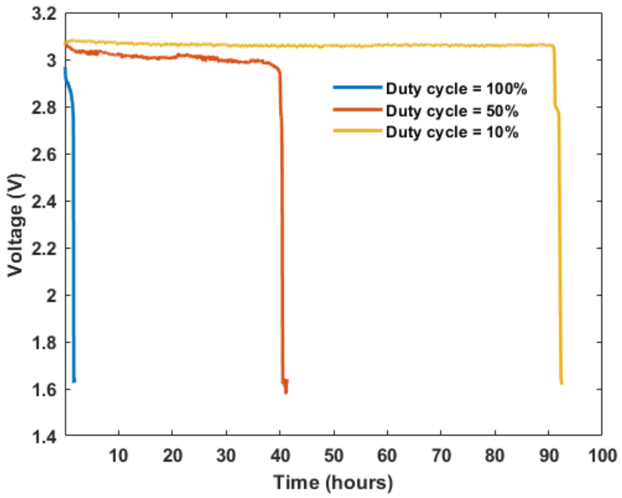

**Figure S8. Measured battery voltage over time under various duty cycles.** Measured battery life of the whole system was evaluated under various duty-cycle configurations, alternating between active and sleep modes. In the sleep mode, the entire system consumes 20 $\mu$A. With 7.5 mAh batteries, a theoretical battery life of up to 14 days can be achieved for duty cycles approaching 1%. In these measurements, battery voltage was sampled every 30 seconds using an analog-to-digital converter (ADC) of the Arduino Uno, which was separately powered by a laptop.

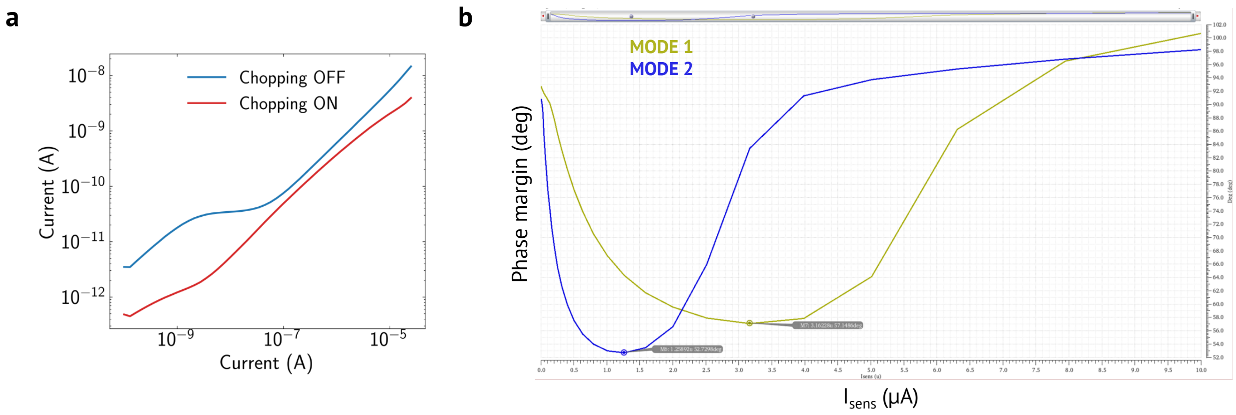

**Figure S9. Simulations of the AFE.**
**(a)** Total integrated noise current of the AFE with chopping enabled and disabled, simulated on Cadence Virtuoso using post-layout parasitic extracted AFE cell. The y-axis represents the total integrated rms noise current from 0.001 Hz to 100 Hz. The x-axis represents the sensor current. As it can be seen, chopping reduces the noise by an order of magnitude at lower sensor currents.
**(b)** Stability analysis that demonstrates the closed-loop phase margin (y-axis) of the AFE under different sensor current values (x-axis), simulated on Cadence Virtuoso using post-layout parasitic extracted AFE cell.

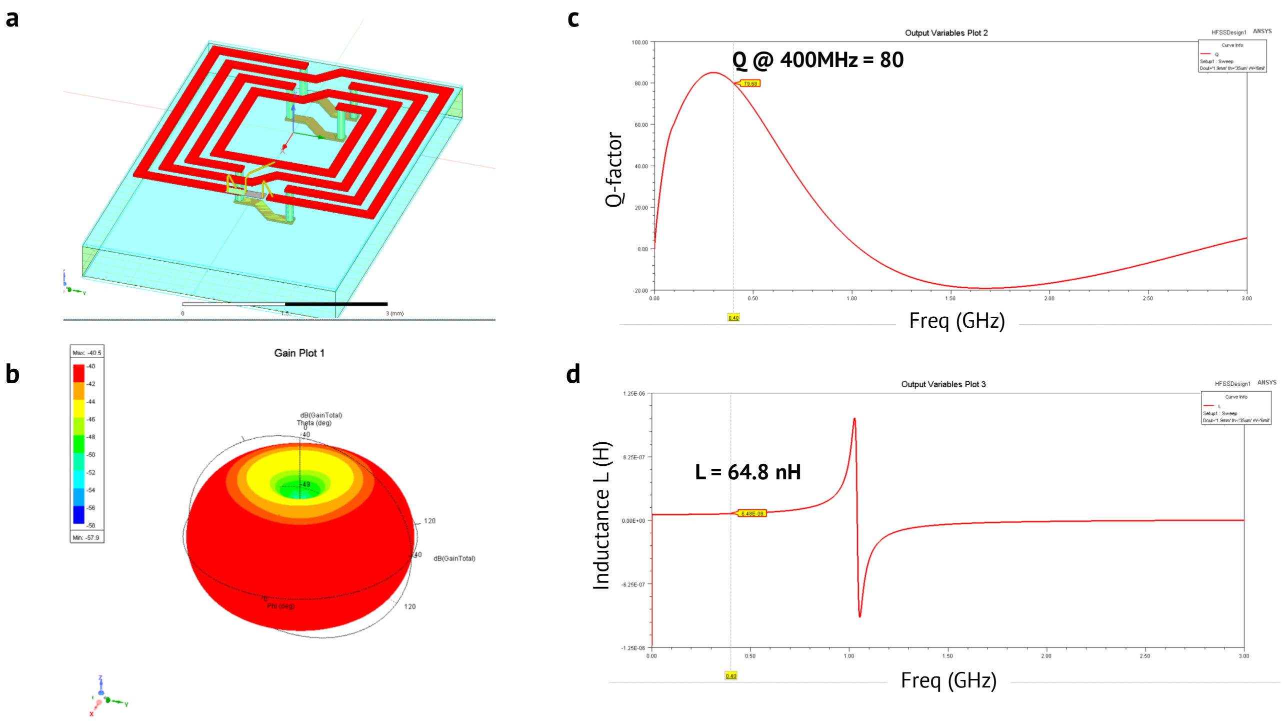

**Figure S10. Simulations of the TX antenna.**
**(a)** The simulation setup of the TX loop coil/antenna on Ansys HFSS using a stack-up of a 4-layer rigid FR-4 PCB. Parameters provided by PCBWay were used.
**(b)** Simulated radiation pattern of the antenna.
**(c-d)** Quality (Q) factor and inductance (L) of the TX coil/antenna.

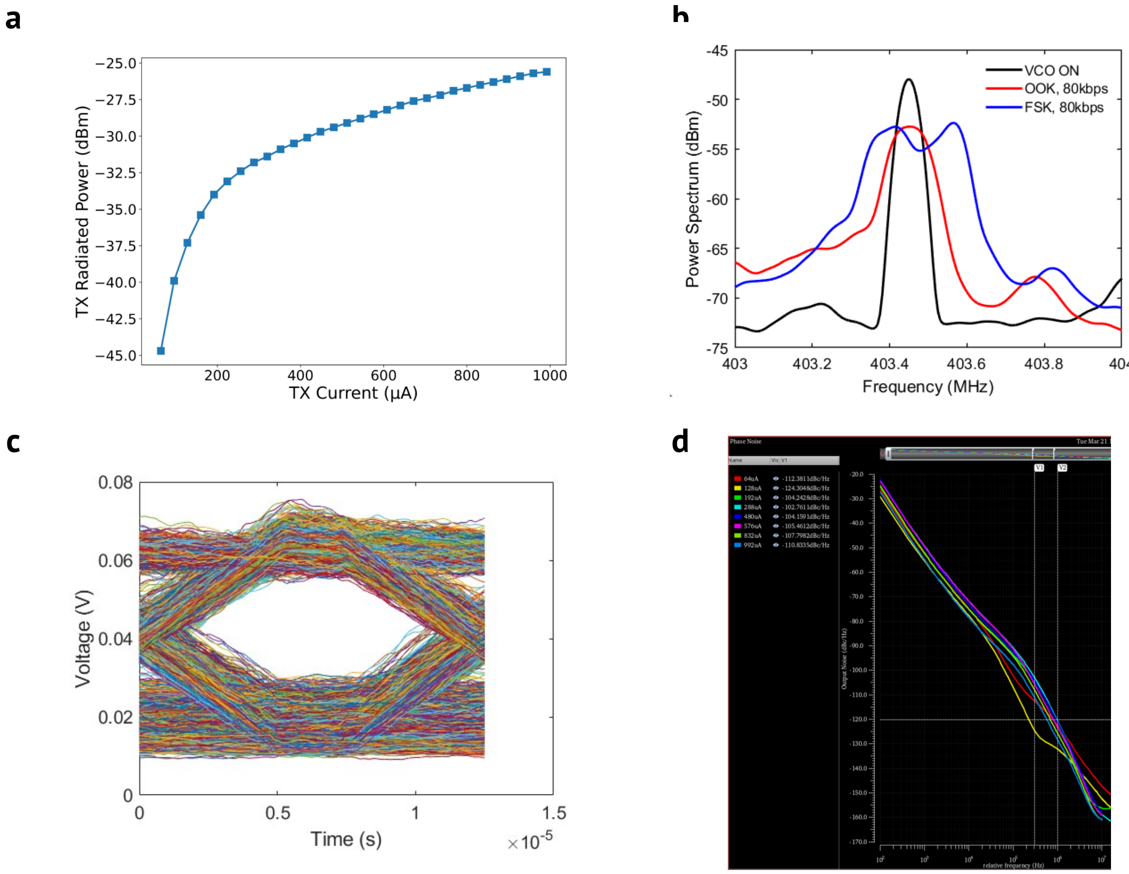

**Figure S11. Measurements and simulations of the TX.**

**(a)** The radiated output power of the TX under various bias currents in the VCO, simulated on Cadence Virtuoso using a post-layout parasitic extracted TX cell and antenna model extracted from Ansys HFSS.

**(b)** Measured frequency spectrum of the TX when using OOK and FSK modulations.

**(c)** Measured eye diagram of the received signal. The TX transmits a PRBS bitstream at 80 kbps, and the external reader is placed 1 m away. The received signal is filtered, and an eye diagram is generated using MATLAB.

**(d)** The phase noise of the TX under various bias currents in the VCO, simulated on Cadence Virtuoso using post-layout parasitic extracted TX cell and antenna model extracted from Ansys HFSS.

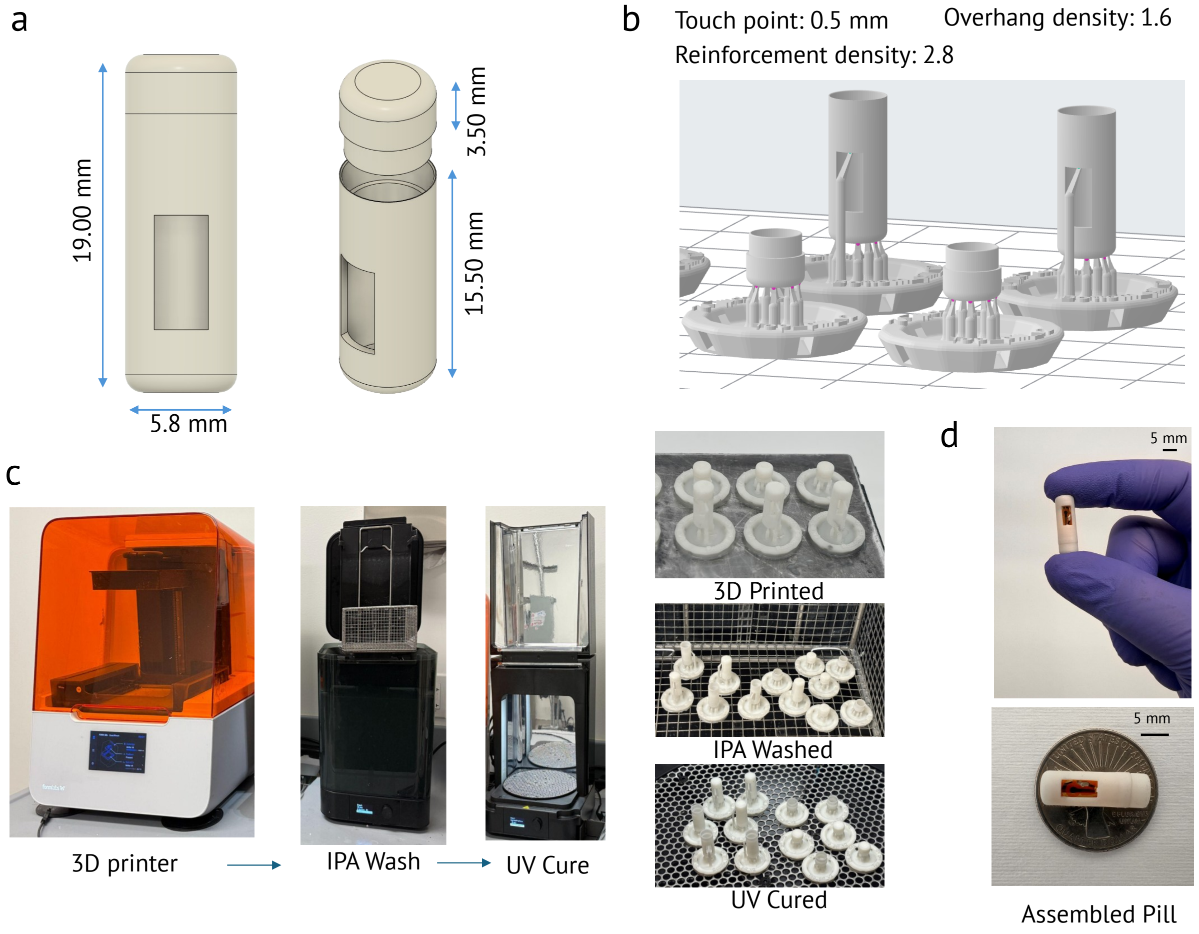

**Figure S12. 3D printing of pill casing.**
**(a)** CAD design of the pill created in Fusion 360, showing key dimensions: 19 mm total length and 5.8 mm diameter, consisting of a 15.5 mm bottom section and a 3.5 mm top cap.
**(b)** Slicing and support parameters used for 3D printing, optimized for the Form 3 printer using biocompatible BioMed White resin.
**(c)** Prototyping workflow including 3D printing, IPA washing for 20 minutes to remove uncured resin, and UV curing for 30 minutes at 60 °C to achieve full polymerization.
**(d)** Final assembled pill integrating the electronics and chemical sensors.

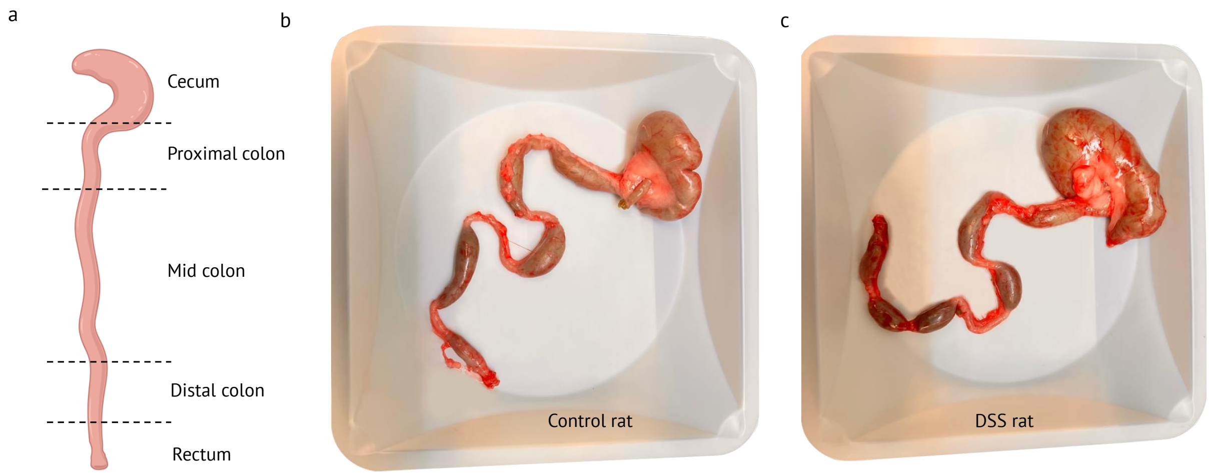

**Figure S13. Colon extraction for histology and ELISA analysis.**
**(a)** Schematic illustration of the different regions of the rat colon, including the cecum, proximal colon, mid colon, distal colon, and rectum. Samples from the distal colon of all six rats were utilized for histology and ELISA analysis.
**(b)** Representative image of the extracted colon from a control rat.
**(c)** Representative image of the extracted colon from a DSS rat, showing visible inflammation induced by acute colitis.

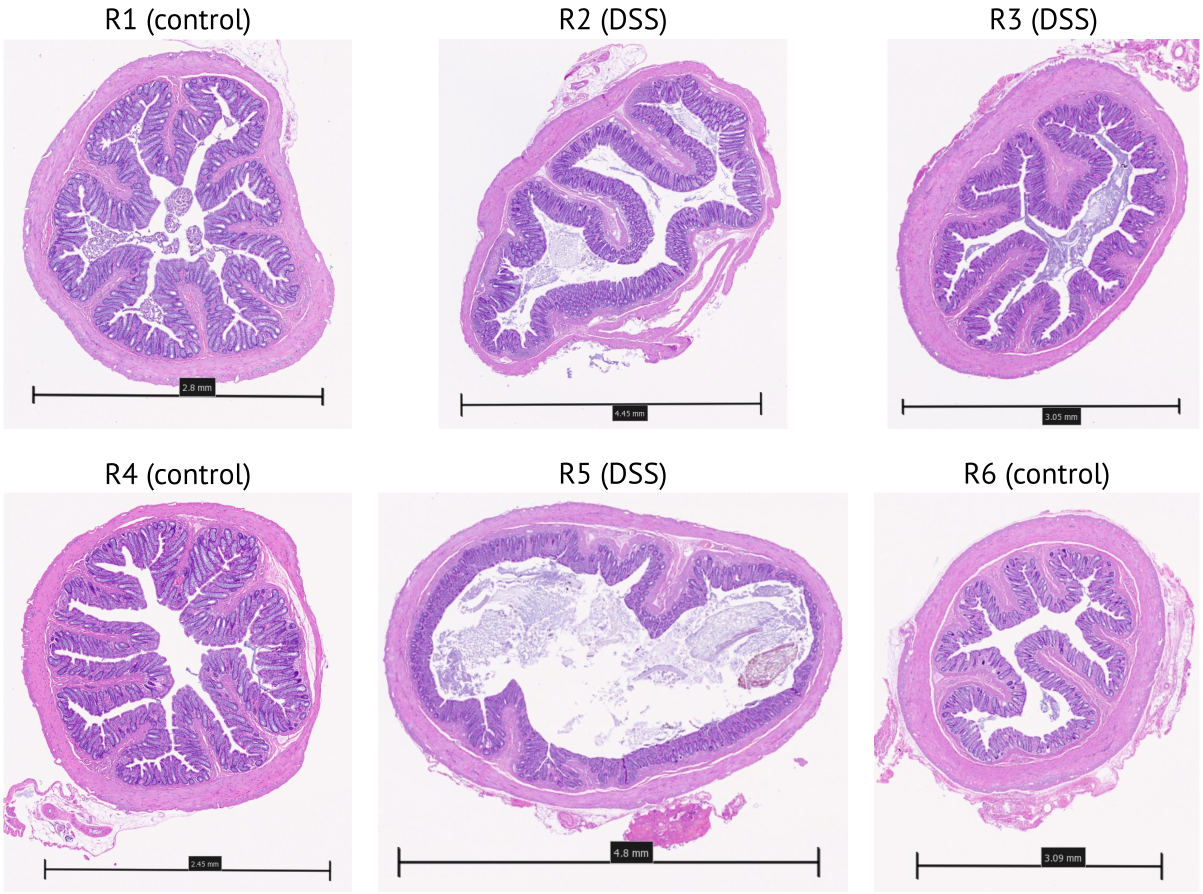

**Figure S14. Histology images.** Histology images of the H&E-stained colon tissues of control and DSS-treated rats after 10 days.

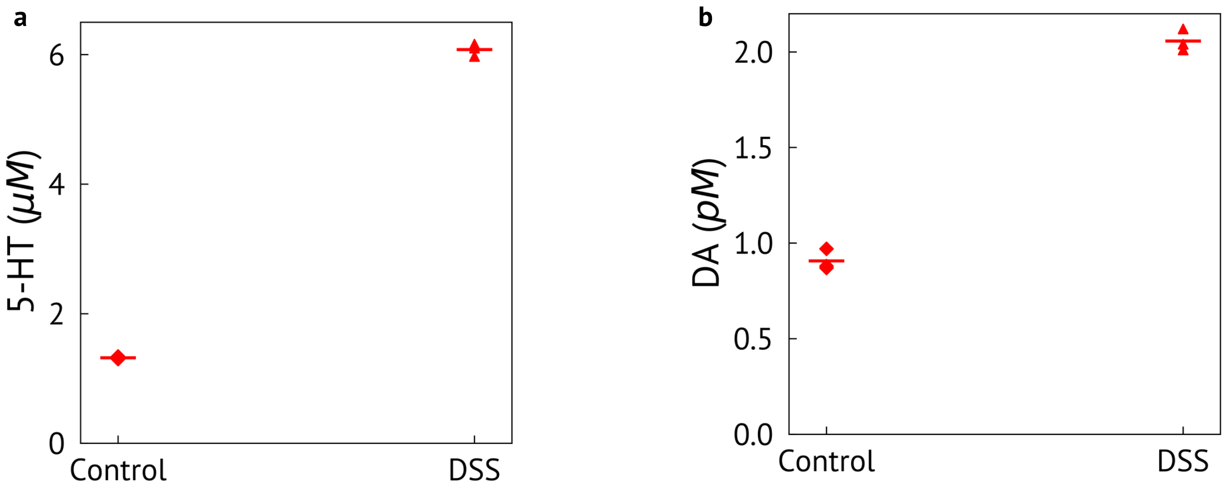

**Figure S15. ELISA analysis.** Rats were administered DSS from day 0 to induce colitis, and measurements were performed on day 10. One rat (R6) from the control group and one rat (R5) from the DSS-treated group were used for ELISA analysis. From each rat three samples were collected from the distal colon to perform the analysis.
**(a)** 5-HT concentrations.
**(b)** DA concentrations.
5-HT concentrations from ELISA analysis were consistent with our sensor data. DA concentrations were negligible in the rat colon, as confirmed by ELISA analysis.

**Table T2.** Comparison with prior ingestible electrochemical sensing systems

| | This work | Min et al.[3] | Even et al.[4] | Xu et al.[5] |
|---|---|---|---|---|
| **Biomarkers** | Serotonin, dopamine | Serotonin, glucose, pH, ionic strength | ORP, pH | pH |
| **Sensing technique** | Amperometry, voltammetry | Amperometry, voltammetry, potentiometry, impedance | Amperometry | Amperometry |
| **Electronics** | Custom IC (AFE, ADC, TX, PMU) & off-the-shelf (MCU) | Off-the-shelf | Off-the-shelf | Custom IC (AFE, ADC, PMU) & off-the-shelf (TX, MCU) |
| **Average current consumption** | 42 $\mu$A$^{\#}$ | 350 $\mu$A$^{*}$ | 28 $\mu$A$^{*}$ | 16 $\mu$A$^{+}$ |
| **Power source** | Battery (1.55 V, 7.5 mAh, ∅ 4.8 mm × 1.6 mm) | Battery (1.55 V, 16 mAh, ∅ 5.8 mm × 2.0 mm) | Battery (1.55 V, 12.5 mAh ∅ 5.8 mm × 1.6 mm) | Battery (N/R) |
| **Pill dimensions** | ∅ 5.8 mm × 19 mm | ∅ 7 mm × 25 mm | ∅ 7.5 mm × 21 mm | ∅ 11 mm × 26 mm |
| **Pill volume** | 502 mm$^3$ | 962 mm$^3$ | 927 mm$^3$ | 2470 mm$^3$ |
| **Experiments** | *In vivo* (rats) | *In vivo* (rats and rabbits) | *In vivo* (pigs and humans) | *In vivo* (humans) |

N/R – not reported, $^{\#}$10% duty cycle, $^{*}$duty cycle values were not reported, $^{+}$20% duty cycle

#### References

1. Shan, B., & Cho, K. (2004). Ab initio study of schottky barriers at metal-nanotube contacts. *Physical Review B—Condensed Matter and Materials Physics*, *70*(23), 233405.
2. Zhong, H., Quhe, R., Wang, Y., Ni, Z., Ye, M., Song, Z., Pan, Y., Yang, J., Yang, L., Lei, M., et al. (2016). Interfacial properties of monolayer and bilayer mos2 contacts with metals: Beyond the energy band calculations. *Scientific reports*, *6*(1), 21786.
3. Min, J., Ahn, H., Lukas, H., Ma, X., Bhansali, R., Sunwoo, S.-H., Wang, C., Xu, Y., Yao, D. R., Kim, G., et al. (2025). Continuous biochemical profiling of the gastrointestinal tract using an integrated smart capsule. *Nature Electronics*, 1–12.
4. Even, A., Minderhoud, R., Torfs, T., Leonardi, F., van Heusden, A., Sijabat, R., Firfilionis, D., Castro Miller, I. D., Rammouz, R., Teichmann, T., et al. (2025). Measurements of redox balance along the gut using a miniaturized ingestible sensor. *Nature Electronics*, *8*(9), 856–870.
5. Xu, F., Yan, G., Wang, Z., & Jiang, P. (2015). Continuous accurate ph measurements of human gi tract using a digital ph-isfet sensor inside a wireless capsule. *Measurement*, *64*, 49–56.
